# Temporal restriction of RNAi reveals breakdown of the segmentation clock is reversible after knock down of primary pair rule genes but not Wnt-signaling in the red flour beetle

**DOI:** 10.1101/2023.10.15.562380

**Authors:** Felix Kaufholz, Julia Ulrich, Muhammad Salim Hakeemi, Gregor Bucher

**Affiliations:** Göttingen Graduate School for Neurosciences, Biophysics, and Molecular Biosciences (GGNB); Johann-Friedrich-Blumenbach-Institut, GZMB, Universität Göttingen, Justus-von-Liebig-Weg 11, 37077, Göttingen, Germany

**Author notes:** Corresponding author: Gregor Bucher, **Email:**. these authors contributed equally (Felix Kaufholz, Julia Ulrich). **Author Contributions:** FK: Investigation, formal data analysis, visualization of the data shown in the main paper; JU: Investigation, formal data analysis, visualization of the establishment of the new tool and of data shown in the supplementary; MSH: Investigation on other uses of the inhibitor; GB: Conceptualization, funding acquisition, supervision, writing the original draft. **Competing Interest Statement:** The authors declare no competing interests. **Classification:** Biological Sciences, Developmental Biology.

**Keywords:** Insect segmentation, clock and wavefront/speed gradient model, segmentation breakdown, RNAi, transgenic tool

## Abstract

Animals from all major clades have evolved a segmented trunk, reflected for instance in the repetitive organization of the human spine or the insect segments. These units emerge during embryonic segmentation from a posterior segment addition zone, where repetitive gene activity is regulated in a spatiotemporal dynamic described by the clock and wavefront/speed gradient model. This model has been tested in the red flour beetle *Tribolium castaneum* and other insects by studying the effect of the RNAi knockdown of segmentation genes. For upstream components such as primary pair rule genes, caudal or Wnt pathway components, this treatment often led to the breakdown of segmentation. However, it has remained untested, how the system would react to a temporally limited interruption of gene function. In order to ask such questions, we established a novel experimental system in *T. castaneum*, which allows blocking an ongoing RNAi effect with temporal control by expressing a viral inhibitor of RNAi. We show that the *T. castaneum* segmentation machinery re-established after we blocked an ongoing RNAi response targeting the primary pair rule genes *Tc-eve, Tc-odd* and *Tc-runt*. However, we observed no rescue after blocking RNAi responses targeting Wnt pathway components. We conclude that the insect segmentation system contains both, robust feedback-loops that can re-establish and labile feedback loops that can breakdown irreversibly. This combination may reconcile two partially conflicting needs of the embryonic regulation of segmentation: A tightly controlled initiation and maintenance of the SAZ by labile feedback-loops ensures that only one segment addition zone is formed. Conversely, robust feedback-loops confer developmental robustness required for proper segmentation, which may be challenged by internal or external disturbances. Our results ponder the insect segmentation machinery from a different angle and introduce a new experimental tool for temporal control on RNAi.

**Significance statement:** The generation of repetitive body parts during embryonic segmentation has been of key interest to developmental biologists, who usually used permanent knock-down of gene function for their studies. Using a new tool to temporally stop a gene knock-down effect, we find both robust and labile feedback-loops within the segmentation machinery. Thereby, the embryo may ensure that only one trunk is formed but that trunk formation is robust against external disturbance.

## Introduction

A striking feature of many animal body plans is their subdivision into repetitive units and clades with segmented bodies, namely vertebrates, annelids and arthopods, are found in all major branches of animal phylogeny (1–3). The repetitive design facilitated the evolution of an amazing morphological and functional diversification along the body axis, contributing to the evolutionary success of these clades. In most vertebrates and arthropods, embryonic segmentation is generated by a posterior clock-like mechanism that uses temporal oscillations of gene activity to generate repetitive spatial patterns (1, 4). Most experiments studying the clock in insects used parental RNAi (5), which leads to knock-down of gene function from the beginning of development. Hence, these experiments revealed only the first essential function of the respective genes and later aspects of the segmentation clocks have remained inaccessible to functional investigations. Here, we present a novel tool for shutting down an ongoing RNAi response with temporal control. We use this method to ask the novel question, whether the segmentation clock can re-establish itself after it has broken down as consequence of a knock-down of key segmentation genes.

In insects, the process of embryonic segmentation has been best studied best in *D. melanogaster melanogaster*, where a hierarchically organized gene regulatory network (GRN) leads to an almost simultaneous formation of all segments (6, 7). In most insects, however, segments are added sequentially from a posterior segment addition zone (SAZ) (1, 1–3). In those animals, a clock-like mechanism seems to sequentially generate the segment boundaries (8–11). Intriguingly, the logic underlying the insect segmentation clock is similar to the one of the vertebrate somitogenesis clock although the involved genes differ (4, 12, 13). The red flour beetle *T. castaneum* has been the main insect model organism for studying the segmentation clock of insects. Current models on its molecular setup have been discussed extensively in recent reviews (1, 4) such that only briefly outline will be given here. In principle, the clock acting in the SAZ consists of two components: A gene or GRN able to oscillate in a cell autonomous way in the SAZ and a posterior to anterior signaling gradient called speed regulation gradient. This gradient across the SAZ remains stable throughout segmentation and activates the cellular oscillator in a concentration dependent way. The combination of both components leads to dynamic on and off states of oscillator gene expression in pseudo-waves initiating in broad domains at the posterior, moving towards the anterior SAZ while becoming narrower and eventually stalling at the anterior boundary of the SAZ to form a new segment boundary. In this work, we use the concept of “speed regulation” instead of the initially suggested “wave front” concept, which actually represents an extreme case of speed regulation (14). In *T. castaneum*, Wnt signalling at the posterior pole of the SAZ is autoregulatory and it activates *Tc-caudal* expression (15–17). A feedback loop between *Tc-caudal* and Wnt signaling has been suggested based on respective spider data and based on the fact that knock-down of both is required for the generation of double head embryos in *T. castaneum* (15, 18). One or both components may function as the molecular realization of the speed regulation gradient (19, 20). *Tc-caudal*, *Tc-dichaete* and *Tc-odd-paired* have been termed timing factors reflecting their subsequent functions in *D. melanogaster* segmentation and their expression in the *T. castaneum* in SAZ in patterns compatible with similar temporal input to the clock (21). The primary pair rule genes (pPRGs) *Tc-even-skipped (Tc-eve)*, *Tc-runt* and *Tc-odd-skipped (Tc-odd)* are the oscillating genes (8, 11) while *Tc-hairy* may have lost an ancestrally essential function in *T. castaneum* (1, 22). Regulatory interactions among the pPRGs are thought to realize the negative feedback loop required for the oscillator. Together, they regulate the expression of the secondary pair rule genes *Tc-paired (Tc-prd)* and *Tc-sloppy-paired*, which eventually turn on the segment polarity genes such as *Tc-wingless (Tc-wg)* (23, 24). The expression of *Tc-ev*e in stripes in the SAZ and *Tc-wg* marking established segment boundaries are shown in Fig. 1A. Different regulatory interactions among the pPRG oscillator genes have been proposed to explain their expression patterns (8). Different from vertebrates, there is another clock ticking in parallel to the periodic pPRG based clock. This non-periodic clock is formed by the *T. castaneum* gap genes and is probably under the control of the same speed regulation gradient. It leads to the one-time sequential activation of the different gap genes (20, 25). While it has become clear that gap genes regulate Hox gene expression for regionalization of the body, they may provide additional input for pPRG regulation as well. Interestingly, the knock-down of several gap gene orthologs led to a complete breakdown of segmentation (22, 26–28).

**Figure 1.**
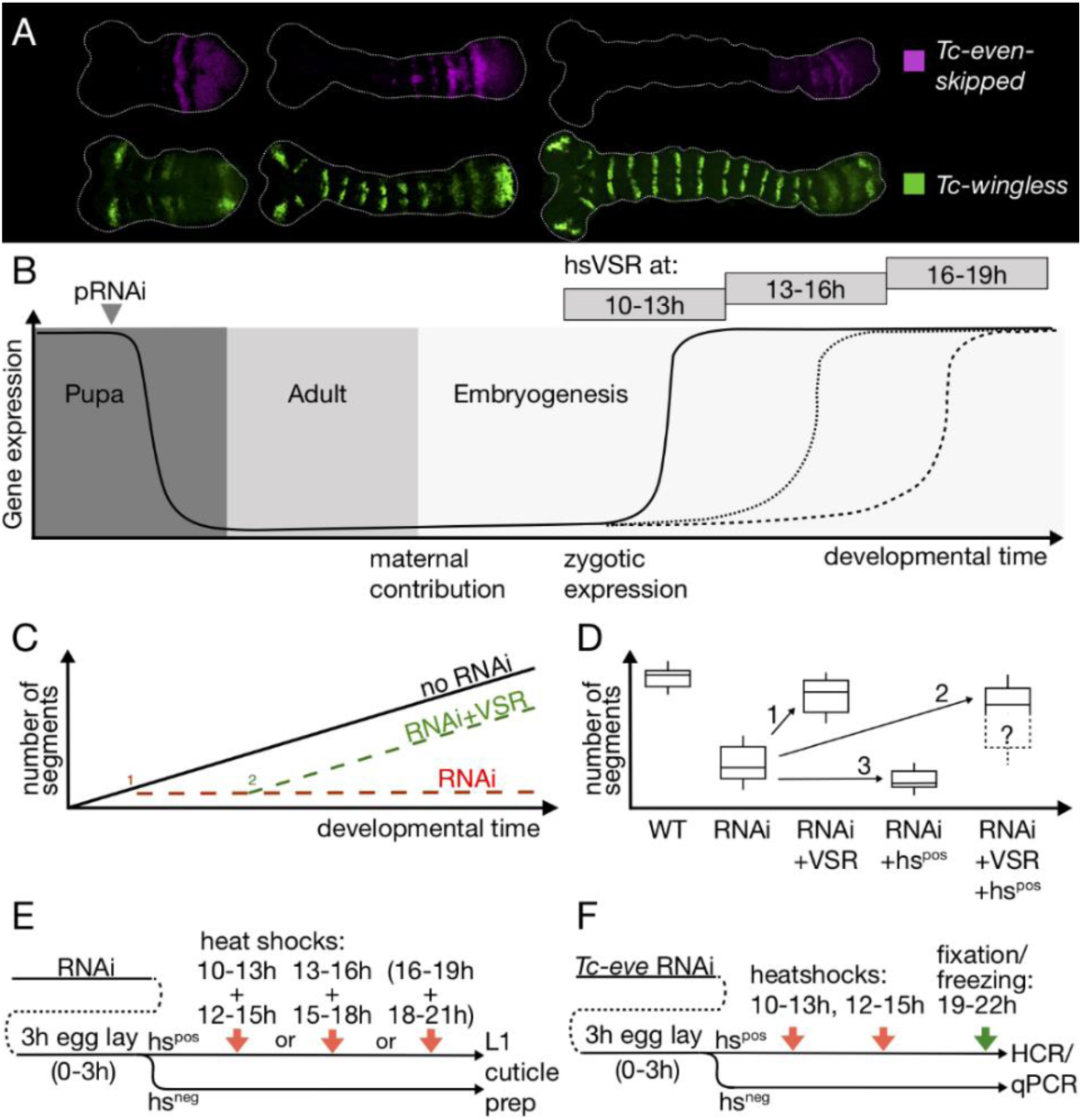
Overview on the experimental design. A) *T. castaneum* embryos representing the stages used for blocking the RNAi (11.5, 14.5 and 17.5 hours of development at 32°C). Shown is the expression of *Tc-eve* as an example for a pPRG and *Tc-wg* as marker for segment boundaries. B) Design of the rescue experiments. After parental RNAi (pRNAi), the level of expression drops (declining black line) and is low in the adult and at the beginning of embryogenesis of the offspring. Heat-shock mediated expression of the VSR blocks the RNAi effect after 10-13, 13-16 or 16-19 hours after egg laying, respectively. This allows gene expression to resume at different developmental stages (increasing black and dotted lines). C) Blocking the studied genes from the beginning of segmentation blocks the formation of any segments (red broken line). In case of heat-shock mediated rescue (timing: 2) segmentation resumes and forms some posterior segments (green broken line). Anterior segments should not be rescued. D) Several additive effects influence the final phenotype. RNAi lowers the number of segments due to the knock-down of an essential patterning gene. Rescue by the VSR increases the number of segments (arrow 1). However, heat-shock as such has negative influence on segmentation making phenotypes stronger (arrow 3). Hence, the final phenotype is a combination of rescue and heat-shock defects (arrow 2). E) Details of the procedure. After parental RNAi, eggs were collected for 3 hours (0–3) and treated with heat-shocks at different times of development in separate experiments: early (10-13h), intermediate (13-16 h) and late (16-19h). To maintain a high level of VSR expression, the heat-shock was repeated after 2 hours, respectively. The latest heat-shock showed minor effects and was not included in all experiments. Hs-negative (hs^neg^) siblings were used as controls. F) For staining the embryos and for the qPCR experiments, the treated embryos were fixed at a stage corresponding to 19-22h of development.

These insights were deduced from modelling, gene expression patterns and from knocking-down gene function by parental RNAi. In the latter experimental approach, dsRNA is injected into female beetles, who transmit the RNAi effect to their offspring, which consequently suffers from the RNAi knock-down from earliest embryonic stages onwards. Hence, strictly spoken, the interactions found in these studies are valid for the first rounds of oscillation while later interactions might differ. Unfortunately, the available techniques did not allow studying the later stages of the clock independently from its initiation. As consequence, it has remained unclear, whether the observed breakdown of segmentation after early knock-down of pPRGs was irreversible or, alternatively, that a continued depletion of gene function was required to induce that drastic phenotype. Similarly, it has remained unclear, whether the observed Wnt autoregulation at the posterior pole was sufficient for maintaining Wnt ligand expression or, alternatively, whether an as yet unknown upstream factor was required for maintaining Wnt signalling at the posterior pole. Finally, it has remained unclear, in how far the network active in the SAZ would be able to re-establish itself after it had broken down.

In order to answer these questions and to open up new experimental possibilities more generally, we developed a system for blocking an ongoing RNAi response with temporal control. For that purpose, we used viral suppressors of RNAi (VSRs), which are proteins that evolved to rescue viruses from the RNAi immune response of the host. We found that heat-shock mediated expression of one VSR, CrPV1A, efficiently blocked the RNAi response in *T. castaneum*. Using this tool, we found that the segmentation breakdown due to pPRG RNAi was reversible, i.e. the system re-established itself once the knock-down was suppressed. In contrast, the breakdown observed after knocking down Wnt signaling components was irreversible. This is evidence that the Wnt autoregulatory loop is at the top of speed regulation gradient maintenance.

## Results

### Establishing a viral suppressor of RNAi as a tool in T. castaneum

RNAi is an anti-viral defense and viruses evolved proteins to interfere with that process. A variety of viral suppressors of RNAi (VSRs) from insect and plant viruses have been described (29–36). Based on the conservation of the proteins involved in RNAi (37) and the proven functionality of some of these inhibitors in flies, we assumed that VSRs might be able to block RNAi in *T. castaneum* as well. To test this, we generated transgenic lines for six VSRs where the VSR expression was under the control of the Gal4 controlled UAS-enhancer (38, 39). These lines were tested for their efficacy in suppressing RNAi in *T. castaneum* by two independent tests. We used two different Gal4 driver lines and tested the rescue of an endogenous gene and a heterologous gene expressed from a transgenic construct (see Supplementary Text 1 for experimental details and results). Only the VSR CrPV1A derived from the *Cricket Paralysis Virus* showed strong reduction of the RNAi effect in both tests while FHV B2 from the *Flock House Virus* showed some effect in one test (see Supplementary Text 1). Hence, we decided to use CrPV1A for our purpose. The CrPV1A protein is responsible for the high pathogenicity of the *Cricket Paralysis Virus* by interacting with the endonuclease Ago-2, a component of the RISC complex (see Supplementary Text 2 for further information). In *D. melanogaster*, it did not interfere with the miRNA pathway (33). However, we were not able to generate transgenic lines with a high level of ubiquitous CrPV1A activity despite many trials. Therefore, we hypothesize that strong ubiquitous VSR expression may affect viability – possibly via blocking the miRNA pathway. Taken together, our results identified CrPV1A as a potent inhibitor of RNAi in *T. castaneum*.

### Temporal control of RNAi by heat-shock mediated VSR expression

In order to gain temporal control on RNAi, we established transgenic lines where CrPV1A was under the control of the *T. castaneum* heat-shock promoter (hsVSR) (40). In order to test the hsVSR for applicability for the segmentation process, we performed several control experiments. As positive control, we chose the secondary pair-rule gene *Tc-paired (Tc-prd),* which is a downstream gene of the segmentation clock (23, 24). Therefore, rescue of segmentation by VSR expression was expected because the segmentation clock does not breakdown in *Tc-prd* RNAi. We performed parental RNAi of *Tc-prd* in our hsVSR line. The RNAi embryos were either not heat-shocked or were treated with heat-shocks during one of three different time windows during germ band elongation (see scheme in Fig. 1B). In the absence of a heat-shock, the L1 larval cuticles displayed the published pair-rule-gene phenotype where the number of abdominal segments was halved to a median of four abdominal segments (Fig 2A,B hs^neg^, J) (23, 24). Heat-shock mediated VSR expression at 10-13 h after egg laying (32°C) rescued the abdominal segment number to a median of 7.5 abdominal segments, i.e. almost to wildtype (Fig 2B, hsVSR 10-13h). Some rescue of more anterior segments was observed as well, i.e. Md (25%), Lab (30%) and the second thoracic segment (80%) (Fig. 2A). Later VSR expression (13-16 h after egg laying) rescued to a median number of only six abdominal segments (Fig 2B, hsVSR 13-16h) while the latest heat-shock (16-19h) failed to rescue abdominal segments (Fig. 2B, hsVSR 16-19h). In our negative controls, i.e. *Tc-prd* RNAi in *vermillion white* (*vw)* wildtype, the cuticles showed no rescue irrespective of whether heat-shock was applied or not (Fig 2A-B). Taken together, this experiment showed that rescue of the segmentation process from an ongoing RNAi effect was possible by blocking RNAi with our hsVSR (see Fig. S1G,H for an independent replicate of this experiment done by another experimenter).

**Figure 2.**
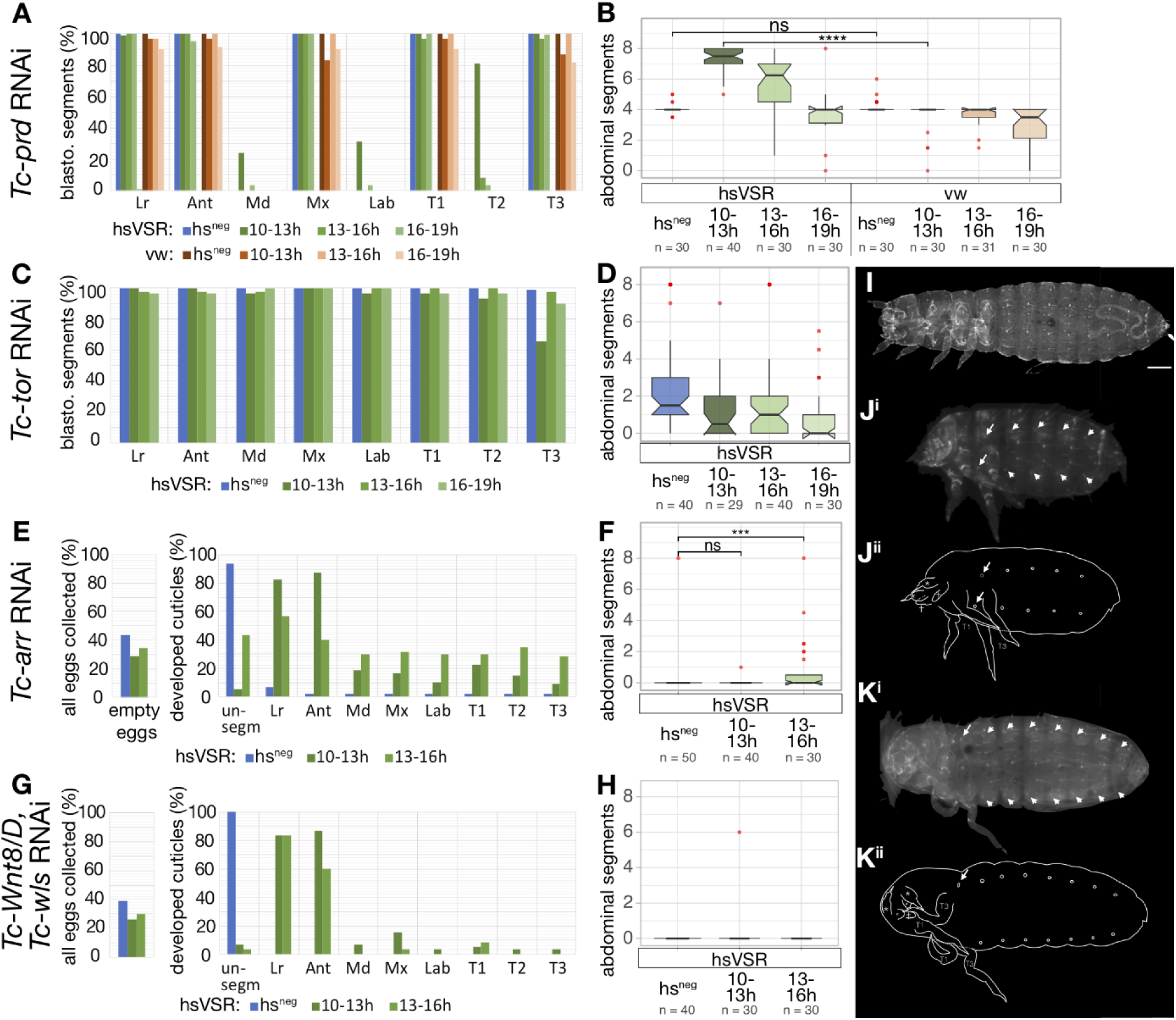
Testing the hsVSR system and rescuing Wnt pathway components. A) *Tc-paired* was used as positive control. The formation of anterior segments, which are built from the blastodermal fate-map, was scored after parental RNAi in the hsVSR line and heat-shock treatments after different times after egg laying (green bars). As negative controls, knock-down embryos of the hsVSR line without heat shock were analyzed (hs^neg^, blue bars) as well as wildtype animals (vermillion^white^ strain, *vw*,) with RNAi and heat-shock (brownish bars). The presence of respective structures was quantified (Lr: labrum; Ant: antanna; Md: mandible; Mx: maxilla; Lab: labium; T1-3: thoracic segments). As expected, Md, Lab and T2 were mostly absent after RNAi. Only the early heatshock treatment in the hsVSR line (10-13h, dark green bars) showed clear rescue. In line with segmentation proceeding from anterior to posterior, the posterior segments were rescued more frequently. B) In the same animals, the number of abdominal segments was counted. In the wt RNAi control (right part, brownish boxes) the number of segments was reduced to four by RNAi as expected. The same was found for the non-heat-shock control of the hsVSR line (leftmost box, hs^neg^). Rescue was observed in the heat-shocked hsVSR animals. It was strongest for the earliest heat-shock (dark green box) while no rescue was observed in the latest (light green box). As unspecific side effect, the heat-shock treatment increased the severity of the RNAi treatment in both hsVSR and wt animals (reduced number of segments seen in right-most green and brownish boxes). C,D) The same treatments were performed for the negative control *Tc-tor*. No rescue was observed as expected for this gene, which is active in the SAZ only at the initiation of segmentation. E-H) Knock-down of Wnt components is known to lead to empty-egg phenotypes, where embryogenesis stops before secretion of a cuticle. Therefore, we documented the portion of empty-egg phenotypes in all collected embryos (left panel in E and G) and analyzed the subset of embryos with cuticle both for the anterior morphological structures (right panel in E) and the number of abdominal segments (F). We found no rescue for Wnt8/wsl double RNAi (G,H). Some rescue of anterior structures and abdominal segments is observed for *Tc-arr* (E). However, we assign this effect to later functions of Wnt, which are independent from its SAZ function (see text for arguments). I,J) In *Tc-prd* RNAi cuticles, loss of anterior and abdominal segments is observed (compare J^i^, J^ii^ to I). K) After heat-shock mediated rescue, the anterior defects are still observed (white arrows) while the posterior abdominal segments are rescued (white arrowheads mark segmental tracheal openings).

As negative control, we tested *Tc-torso (Tc-tor).* In *T. castaneum*, Torso signaling is active at the posterior pole in early embryos but not during elongation. Hence, it was suggested to be required for establishment of the SAZ and to initiate posterior elongation but was unlikely to be required for maintaining it (41). Therefore, blocking RNAi targeting Torso signaling during elongation should not have an effect on posterior segmentation. In line with previous results, our RNAi experiments targeting of *Tc-tor* resulted in the loss of most abdominal segments with a median of 1.5 abdominal segments remaining (blue box in Fig. 2D) while the anterior segments remained unaffected (Fig. 2C) (41). As predicted, no rescue of abdominal segments was observed by hs-induced VSR expression for neither time point (green boxes in Fig. 2D).

Finally, to test for unspecific effects induced by the hs-treatment, we performed the same experiments in the hsVSR line and the vw wildtype strain, which is the genetic background for the hsVSR line. Indeed, the heat-shock treatment alone led to some reduction of abdominal segments. This was obvious in heat-shocked animals of wildtype (Fig. 2B, light red boxes) and in of hsVSR animals without previous RNAi treatment (Fig. S1F). In line with this apparently non-specific effect of heat-shocks and/or VSR expression, the RNAi defects increased in heat-shocked animals in both, *Tc-tor* (Fig. 2D – compare green to blue boxes) and *Tc-paired* knock-down embryos (Fig. S1H). In conclusion, the overall effects observed after RNAi and heat-shock induced rescue is composed of three additive effects: First, the reduction of segments resulting from the specific RNAi effect (Fig. 1D – second box). Second, additional reduction due to unspecific heat-shock and/or VSR defects (Fig. 1D arrow 3) and third, rescue by blocking the RNAi effect by VSR expression (Fig. 1D, arrow 1). The observed overall phenotype results from the combination of these partially opposing effects (Fig. 1D arrow 2).

In summary, these proof of principle experiments showed that the hsVSR system was able to inhibit an ongoing RNAi response where the 10-13 h time window (and to lesser degree the 13-16 h window) appeared optimal for effects on the segmentation process. Further, they revealed side effects induced by the heat-shock treatment.

### Interfering with Wnt signaling leads to an irreversible segmentation breakdown

Parental RNAi targeting several segmentation genes led to the loss of all posterior segments, indicating a breakdown of the segmentation machinery. This phenotype is observed for some gap gene orthologs, primary pair rule genes, the terminal gene torso and two components of the segment addition zone (SAZ), namely caudal and Wnt signaling (8, 26–28, 41–43). In all these experiments, the genes were knocked down throughout development by parental RNAi. Therefore, it has remained unclear, whether the phenotype reflected an irreversible breakdown of segmentation or whether continued depletion of the respective component was required for the continuous loss of posterior segments.

Our new system allowed us for the first time asking whether the segmentation breakdown observed in those RNAi experiments was reversible or not. Wnt signaling and *Tc-caudal* expression are found in the SAZ throughout elongation and respective RNAi experiments led to a segmentation breakdown (42, 43). It was suggested that Wnt regulates *Tc-caudal*, which in turn represents a speed regulation gradient, which is required to regulate the segmentation clock acting in the SAZ. (4, 19). Indeed, autoregulation of Wnt signaling and activation of *Tc-caudal* by Wnt signaling was shown previously for *T. castaneum* (17, 19). It should be noted that in more basal insects and a spider, interfering with Wnt signaling had similar drastic effects on segmentation but the Wnt-cad interactions suggested above were not fully confirmed there (44). At least in *T. castaneum*, an autoregulatory loop is suggested to ensure the continuous expression of these components in the SAZ. Hence, interrupting the loop could to lead to an irreversible breakdown. Alternatively, if a so far unknown upstream signal located in the posterior SAZ was required for their maintenance, the system would be able to re-establish itself. In order to distinguish between these possibilities, we analyzed *Tc-WntD/8,* which together with the Wnt pathway component *Tc-wntless* (*Tc-wls)* is required for posterior segmentation, which is also true for the Wnt receptor *Tc-arrow (Tc-arr)* (42, 45). Besides that role in the SAZ, Wnt is also required for later aspects of segmentation such as the formation of parasegment boundaries. Therefore, the RNAi phenotypes are a mix of early and late functions, which needs to be considered when interpreting the rescue effect. In line with published results, our *Tc-WntD/8+Tc-wls* double RNAi and *Tc-arrow* single RNAi resulted in two classes of phenotypes: Completely unsegmented cuticles and “empty egg phenotypes” (ee-phenotype). The ee-phenotype describes eggshells that do not contain embryonic cuticle, because the embryos stopped development before secreting cuticle while the former are a combination of early and late segmentation defects. We found roughly 40-45% empty eggs for both RNAi treatments (Fig. 2E,G, blue bar in left panel). Most cuticles showed an unsegmented phenotype (90-100%) (blue bars in “unsegm” column of right panel in Fig. 2E and G). hsVSR expression at 10-13h slightly reduced the portion of the ee-phenotype (Fig. 2E and G, green bars in left panels). Strikingly, the portion of unsegmented cuticles dropped dramatically (to roughly 5% for both RNAi treatments, see Fig. 2E and G, compare green to blue bars). The anterior pre-gnathal structures (labrum and antennae) were rescued more strongly than gnathal and thoracic segments. However, no rescue of abdominal segments was observed for the early treatment (Fig. 2F and H; 10-13h). Later VSR expression (13-16h), showed a similar result (Fig 2. E,G; light green bars) but we observed some cuticles with an increased number of abdominal segments after late VSR treatment in the *Tc-arr* RNAi but not *Tc-WntD/8;Tc-wls* RNAi (Fig. 2F, 13-16h). The lack of posterior rescue cannot be due to failed RNAi inhibition, because the clear anterior rescue shows effective inhibition of RNAi by our hsVSR (see Fig. S2 for a biological replicate of both experiments done by another experimenter with similar results). We ascribe the minor posterior rescue seen in the *Tc-arr* experiments (Fig. 2E,F) to the mentioned later Wnt functions (e.g. the formation of segment boundaries) for two reasons: First, the rescue effect increased with the later heat-shocks. This is in contrast to the expectation for upstream components of the SAZ, where a later rescue should only be able to rescue the most posterior segments as was observed in our positive control *Tc-prd* (Fig. 2A,B). Second, those structures, which form independently of the SAZ but need other aspects of Wnt signaling (labrum and antenna) are rescued to a high degree. Third, we do not see rescue when targeting Wnt8, which is exclusively expressed in the SAZ (46).

In summary, our analyses indicated that the breakdown of abdominal segmentation after loss of Wnt signaling was irreversible, indicating the interruption of an essential autoregulatory loop of Wnt signaling alone or autoregulatory interactions between Wnt-caudal. We were not able to test *Tc-caudal* because parental RNAi leads to sterility prohibiting the collection of the high number of embryos required for these type of experiments.

### Segmentation breakdown after primary pair rule gene knock-down is reversible

Downstream of the Wnt signaling- and *Tc-caudal* gradients, three primary pair-rule genes (pPRG) are essential for segmentation. *Tc-even-skipped* (*Tc-eve*), *Tc-runt* (*Tc-run*), and *Tc-odd-skipped* (*Tc-odd*) form a regulatory circuit leading to their oscillating expression in the SAZ. The resulting overlapping stripes provide spatial information for segmentation (10, 11; models discussed in 20, 1, 8, 4). RNAi knock-down of each pPRG leads to the breakdown of blastodermal and posterior segmentation (8). It was suggested that their mutual regulation represented a regulatory circuit, which had to be started at the blastoderm stage and stopped after elongation was completed (8). Later, it was suggested that their oscillations were under the control of a speed regulation gradient provided by ongoing expression of *Tc-caudal* in the SAZ (10, 19, 20). In a scenario with a fully autonomous regulatory circuit, the breakdown would be irreversible while in the model involving a speed regulation gradient, re-establishment of segmentation under the control of the unaffected upstream Wnt/*Tc-caudal* function was likely. As previously shown, *Tc-eve* RNAi resulted in cuticles that retained only labrum (Lr) and antennae (Ant) in both the hsVSR line and the wild type controls (Fig. 3A and C, “hs^neg^”). In contrast to previous results, we noted a pair of tracheal openings (90%, not shown). Expression of the VSR during the early time window (10-13h) did rescue both some anterior and abdominal segments (Fig. 3C and D, green bars and boxes). Rescue of Md, Mx and one thoracic segment was observed (probably T1 as judged by the absence of a tracheal opening) (Fig. 3C). The median number of abdominal segments increased to four segments (with some cuticles showing as many as 6-7 abdominal segments, see Fig. 3D). The hs-treated wild type control did not show any rescue (Fig. 3C,D; red bars and boxes). VSR expression at 13-16h led to no significant rescue. However, some cuticles actually had more than the expected number of abdominal segments but this apparent rescue was counterbalanced by cuticles with additional loss of segments (Fig. 3D, “hsVSR, 13-16”). Likewise, the vw controls showed additional loss of segments upon heat-shock. Hence, it is possible that the negative effect of the hs-treatment counterbalanced a minor rescue effect at the late time window (see Fig. 1D). This experiment was repeated two more times, where one experiment showed similar results and one revealed no rescue effects (Fig. S3).

**Figure 3.**
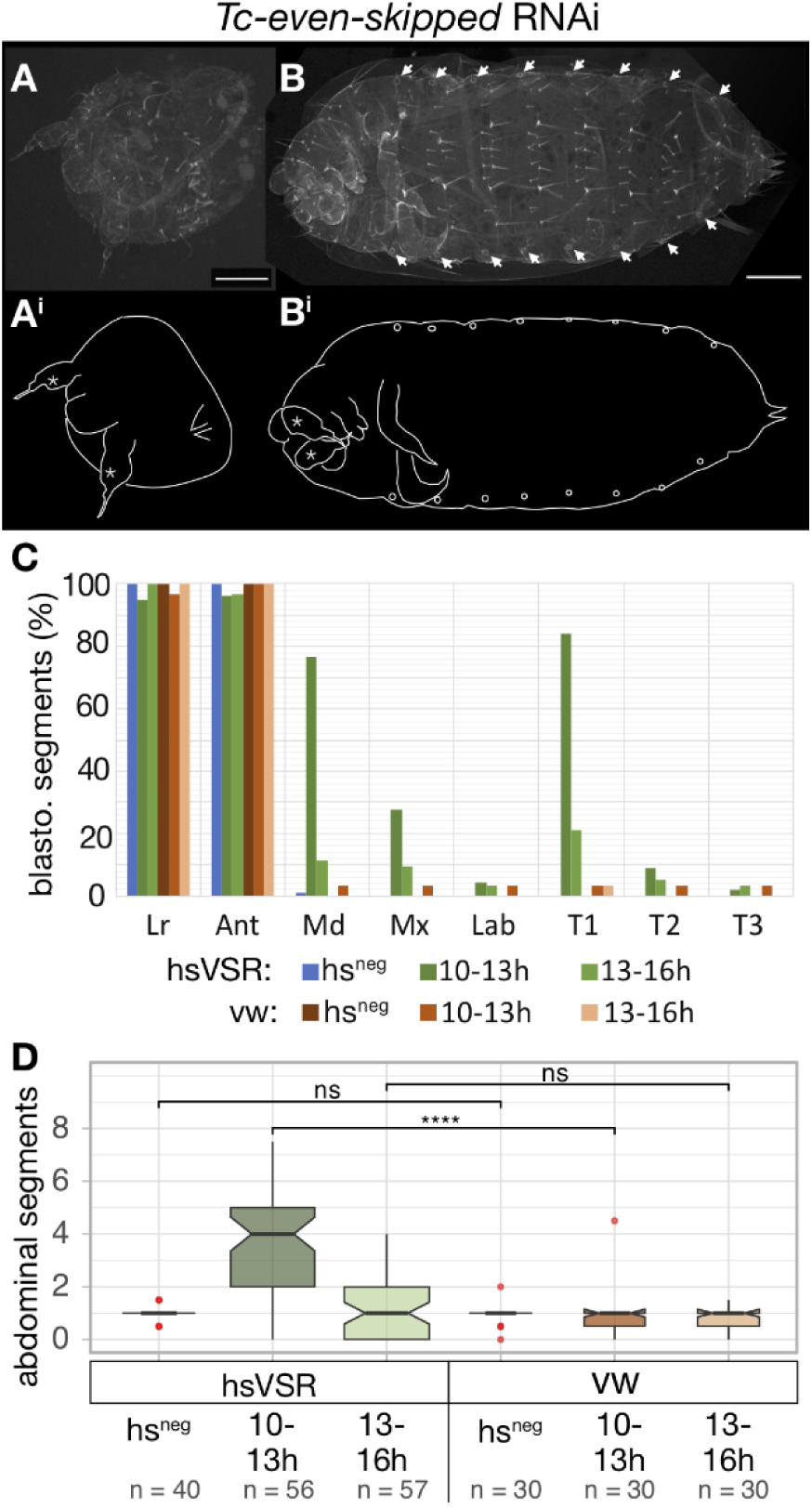
Re-establishment of segmentation after rescue of *Tc-eve* expression. A) *Tc-eve* RNAi leads to complete loss of trunk segmentation – only the labrum, the antennae and the terminal urogomphi can still be discerned in the resulting cuticle balls. B) After hsVSR rescue, some anterior segments in addition to some posterior abdominal segments are rescued – in the specimen shown, all eight abdominal segments are discernable. C) Quantification of the effects for the blastodermal segments reveals the highest degree of rescue for the early treatment. D) Likewise, the most clear rescue of abdominal segments is found for the early rescue (dark green; 10-13h). Labelling as in Fig. 2

As previously published, parental *Tc-runt* RNAi resulted in cuticles that carried mandibles and up to one abdominal segment (Fig. 4A,C). Early VSR expression (10-13h) rescued some blastodermal segments (mx, T1, T2) and the abdominal segments to a median number of three (Fig. 4B, C and D). Some cuticles showed five or more rescued abdominal segments (Fig. 4D). Rescue at 13-16 h showed a similar albeit weaker rescue. Two more repetitions by the same researcher revealed no effect while a repetition by another researcher confirmed the rescue (Fig. S4).

**Figure 4.**
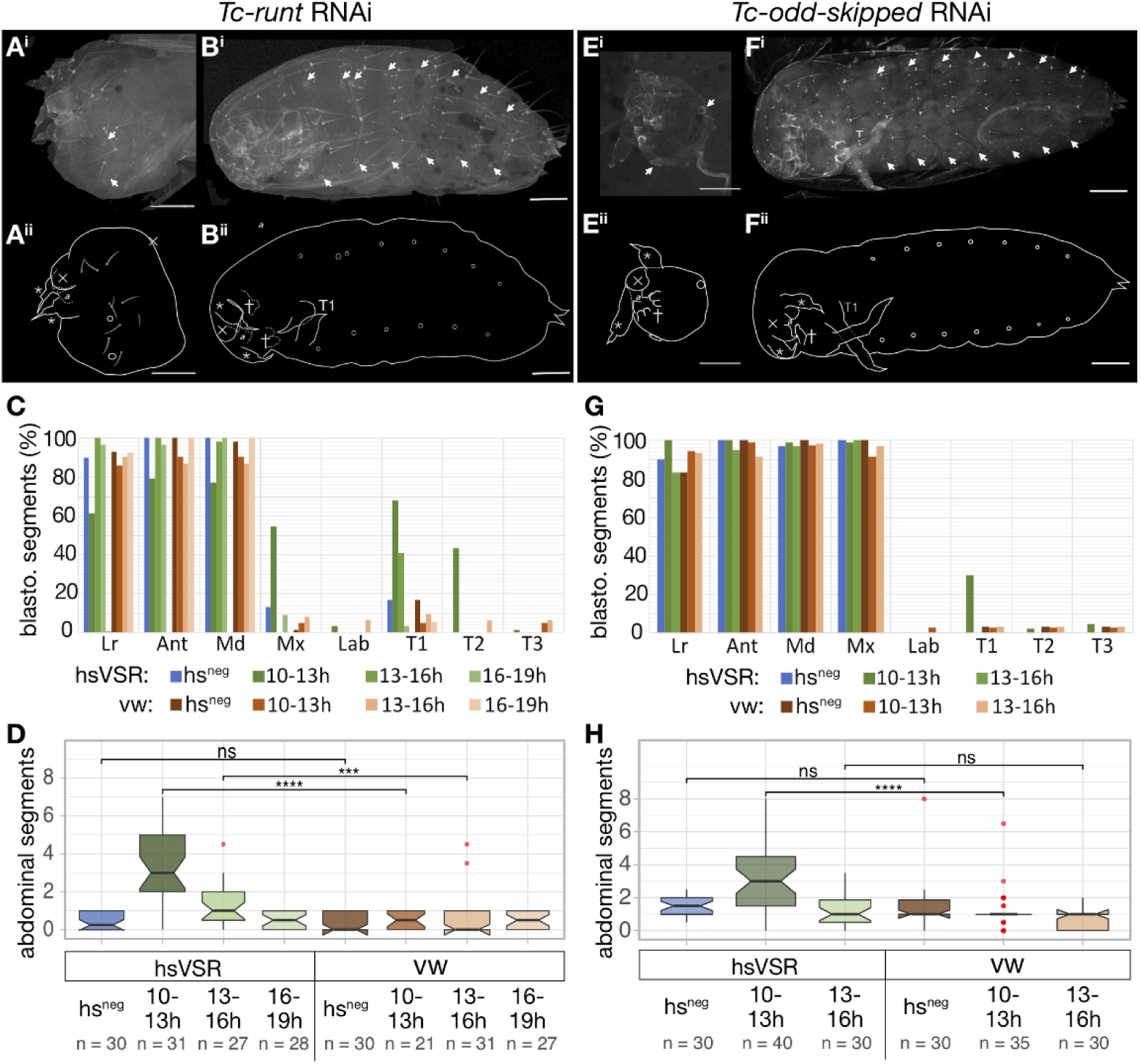
Re-establishment of segmentation after rescue of *Tc-runt* and *Tc-odd* expression. A,B) The *Tc-runt* RNAi phenotype (A) is rescued after the hs-treatment (B). C,D) Quantification of the effects for the blastodermal (C) and the abdominal segments (D) reveals the highest degree of rescue for the early treatment. E,F) Cuticles of *Tc-odd* RNAi phenotypes without (E) and with hsVSR rescue (F). G,H) The rescue of anterior segments (G) and abdominal segments (H) was quantified. The rescue of blastodermal segments is weaker in *Tc-runt* and *Tc-odd* compared to *Tc-eve*. Labelling as in Fig. 2

*Tc-odd* parental RNAi knockdown resulted in cuticles missing all segments posterior to the maxillae (Mx) (Fig. 4E), as expected. Only the early window of VSR expression (10-13h) significantly rescued abdominal segments to a median number of three segments (Fig 4F,H). Of note, a small number of cuticles showed up to 8 rescued segments. Again, two more repetitions gave unclear results while the repetition by another scientist showed a clear effect (see Fig S4).

In summary, for all three pPRG we found that segmentation could be re-initiated after a breakdown. Interestingly, the rescue for the primary pair rule genes was mostly restricted to the earliest time window of RNAi suppression while the rescue of the secondary PRG *Tc-prd* was found also for later time windows. Due to the complex setup, the strict timing requirements and likely variability in the heat-shocks of these experiments, not all experiments led to rescue. However, for each pPRG we found rescue of segmentation in at least two independent replicates and the combination of posterior rescue with anterior deletions is a very specific and unique phenotype. Together with the expression analysis presented below, this makes us confident that the results are valid.

### Gene expression patterns reflect early rescue by hsVSR treatment

Our results on the cuticle level indicated that the segmentation machinery could be re-established after breakdown. However, cuticle is secreted at the end of embryonic development (which takes roughly 72h at 32°C) while segmentation takes place during the first 24 h. Hence, late compensatory effects could blur the early direct effects of the rescue. Therefore, we wanted to observe the re-establishment of the segmentation clock more directly after VSR expression. *Tc-eve* RNAi was chosen for that purpose because it had shown the most robust response in our previous experiments. We repeated the experiment and checked a portion of the embryos for successful rescue on the cuticle level to confirm successful performance of the experiment. The other embryos were fixed some hours after the hsVSR treatment to visualize the expression of the three pPRGs and the segmental marker *Tc-wingless (Tc-wg)* (see Fig. 1F for experimental outline). Of note, we had to fix wt and heat-shocked embryos at different times in order to obtain comparable stages because heatshock leads to a delay in development for which we had to compensate. Hence, we first carefully staged *Tc-wg* patterns (Fig. S5) and optimized the timing of fixation such that animals from the different experimental groups (with and without heat-shock, with and without *Tc-eve* RNAi) would be at comparable stages (Fig. S6).

Parental RNAi of *Tc-eve* in wildtype resulted in the almost complete loss of segmentation irrespective of heat-shock treatment. Instead of segmental stripes, a broad *Tc-wg* domain was observed in the trunk (Fig. 5Aiv and Biv). *Tc-eve* formed one broad domain without stripes (Fig. 5Aiv and Biv), which did not overlap with the *Tc-wg* pattern. Likewise, *Tc-odd* was expressed in one domain while *Tc-runt* showed two abnormal stripes (Fig. 5Aii and Bii – compare to wildtype patterns in C). *Tc-eve* RNAi in the hs-VSR line without heat-shock led to essentially the same patterns (compare Fig 5C to A and B). However, upon hsVSR treatment, the expression of *Tc-wg* reflected the formation of stripes (Fig. 5 Div – compare to Aiv, Biv and Civ). In addition, all three pPRGs re-gained some degree of striped expression (Fig. 5D). To quantify this rescue, we counted the number of stripes of the pPRGs and *Tc-wg* in a number of embryos. Indeed, we found a highly significant increase of stripes after rescue for *Tc-wg* while the low number of pPRG stripes visible at the same time (maximum three) led to a low level of statistical significance (compare the values after heat-shock in Fig. 6A). In an alternative approach, we sorted the stained germband embryos into three classes based on their overall expression patterns: “Close to WT” (WT), “intermediate” (+/-) and “all stripes lost” (-). (Fig. 6B; see Fig. S7 for documentation of pictures and our embryo classification). For all three pPRGs, the highest portion of WT and intermediate phenotypes was found in the hsVSR treated batches (Fig. 6B hs-VSR, 10-13 h). For *Tc-eve*, the difference was statistically significant while for the other genes, the p-value was low but did not reach significance levels (Fig. 6B).

**Figure 5.**
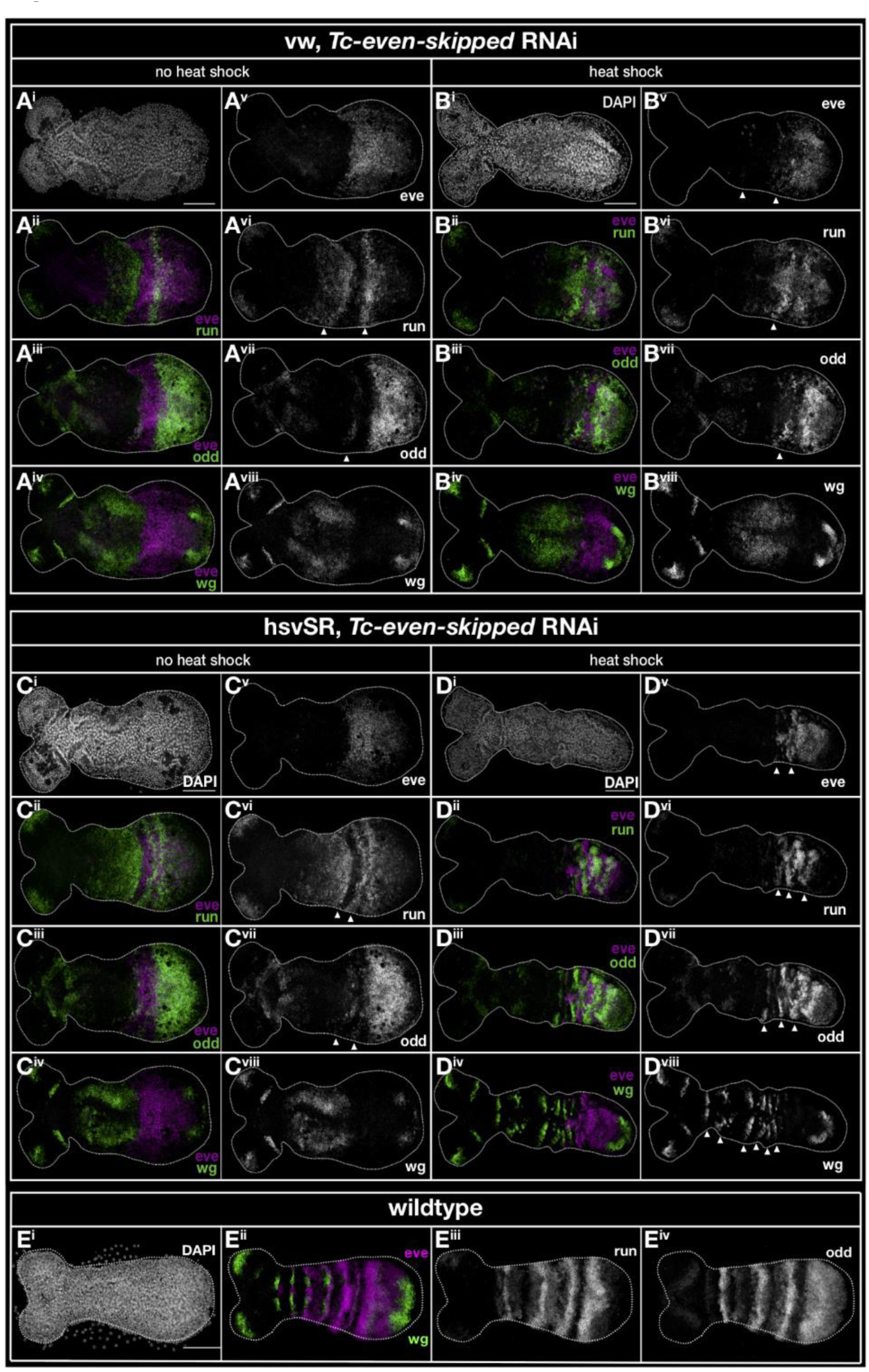
Expression of pPRGs and Tc-wg in Tc-eve RNAi embryos with and without hsVSR rescue. A,B) The expression of the primary pair rule genes and *Tc-wg* are severed in wildtype embryos after *Tc-eve* RNAi (A). Heat-shock alone does not rescue the defects (B). The morphology of the embryo is shown by DAPI staining (A^i^ and B^i^). The quadruple HCR in situ staining in those embryos is shown in combinations of two genes, respectively (left column) and for each gene separately as greyscale picture (right column, respectively). C,D) RNAi in the hsVSR line leads to defects comparable to wildtype (C, compare with A or B). However, hs-treatment does lead emergence of stripes (D). E) The expression patterns are shown in wildtype without RNAi treatment.

**Fig. 6.**
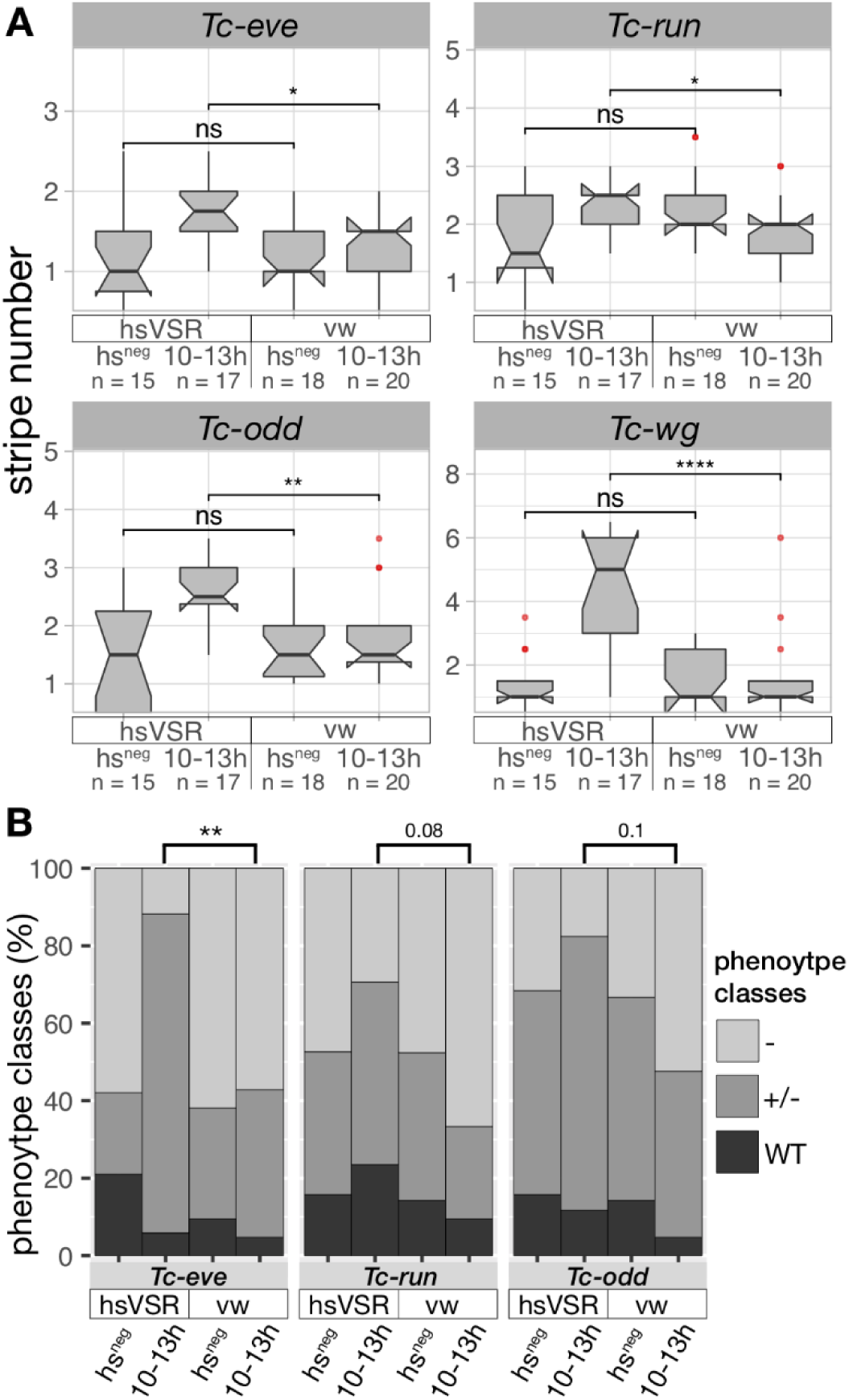
Quantification of gain of striped pPRG expression after hsVSR treatment. A) The number of pair rule gene and *Tc-wg* stripes increased significantly when comparing the heat-shocked batches from the hsVSR line and wildtype. B) In an alternative analysis, the resulting embryos were assigned to three classes: no stripes (-), intermediate (+/-) and close to wildtype (WT). The p-value for Tc-eve reached significance levels while for the other genes the p-value was low but not significant.

### Self-repressing function of Tc-eve revealed by qPCR

Finally, we sought to check for up- and downregulation of the involved genes by qPCR. We confirmed strong increase of VSR expression upon heat-shock (Fig. S8). Expression of *Tc-odd* and *Tc-runt* were not much altered in line with our expression analysis, where both genes remain expressed but lose their striped patterns (Fig. 5). Surprisingly, RNAi targeting *Tc-eve* did not reduce the *Tc-eve* transcript level. However, when testing for intronic sequences in *Tc-eve* RNAi embryos, we found a strong upregulation of expression (Fig. S8). Apparently, the loss of *Tc-eve* function leads to upregulation of its expression, blurring the qPCR results. This result is strong indication for a so far ignored self-repressing function of *Tc-eve* during segmentation. In line with this assumption, *Tc-eve* RNAi embryos that were rescued with hsVSR returned to normal intronic expression levels (Fig. S8).

## Discussion

### Gaining temporal control on RNAi by hsVSR – new possibilities and restrictions

With this work we expand the toolkit of *T. castaneum* with a system that allows to block RNAi with temporal control. The tool might be useful to distinguish between early and late function of genes (shown in this study) or for the analysis of other temporal processes such as the sequential expression of neuroblast timing factors. Further, it could help to overcome technical problems with parental RNAi of genes that lead to sterility. To that end, blocking RNAi in the mother but not the offspring could reduce the sterility issue. We note that the heat-shock experiments were sensitive to changes in the procedure and had to be optimized carefully. Importantly, we show that the negative effect of a heat-shock treatment on developmental processes has to be controlled for. Given the documented function of CrPV1A in flies and beetles, it seems likely that it will be active in other insects, too, opening the possibility to transfer that technique to other species. Theoretically, the CrPV1A VSR could also be used for tools that allow spatial control of RNAi. However, from several unsuccessful attempts in that direction we conclude that strong ubiquitous expression of the CrPV1A probably has negative effects on viability. This may interfere with the establishment and maintenance of lines with strong ubiquitous VSR effect (Hakeemi, in preparation). Using the Gal4/UAS binary expression system for establishing spatial control may be a viable alternative (39).

Of course, it would have been valuable to study the late interactions of the segmentation machinery such as taking out a component during ongoing segmentation. We thought this could be done by dsRNA injection into embryos at different stages. Indeed, we tried several injection time series using *Tc-prd* as target gene. In all batches analyzed, the RNAi knockdown was similar in strength for anterior and posterior segments (i.e. gnathal vs. abdominal segments). The later we injected, the weaker was the phenotype for all body regions alike. Apparently the build up of the RNAi response takes longer than the segmentation process, which takes around 15 hours at (32°C) starting at the differentiated blastoderm stage. This would explain why all segments were affected by the same degree of RNAi knock-down. Other experimental approaches would be needed for taking out a component during elongation.

### Segmentation relies on both, robust and interruptible regulatory feedback loops

Parental RNAi targeting a number of segmentation genes leads to the breakdown of segmentation and loss of all or most trunk segments. This breakdown phenotype was observed after the knock-down of several classes of genes such as the gap-gene orthologs *Tc-hunchback, Tc-Krüppel* and *Tc-giant* (26–28), the primary pair rule genes *Tc-eve, Tc-odd* and *Tc-runt* (8), the posterior marker *Tc-caudal* (43) and components of the Wnt signaling pathway (42, 45) and torso signaling components (41). In all these cases, the phenotypes were experimentally generated by a continuous knock-down of the respective gene function throughout development by an ongoing RNAi response. Hence, two alternative explanations for the breakdown phenotypes remained possible: On one hand, they could reflect an inherent instability of the system that irreversibly breaks down after the removal of an essential component – the continuous knock-down would not have been required for the loss of most posterior segments. Alternatively, the system could be robust and able to re-initiate but that the continuous depletion of an essential component led to the continued interference with segmentation leading to the apparent breakdown phenotype. The results presented here indicate that the gene regulatory system of segmentation actually contains both, a robust down-stream component able to re-initiate and a less robust upstream component that can breakdown irreversibly.

Recent elaborations of the clock and speed regulation gradient model contain two genetic feedback loops. First, a positive feedback loop between Wnt signaling and *Tc-cad* expression, which maintains their continued expression in the SAZ (1, 4). The resulting graded activity represents the speed regulation gradient, which influences the velocity of the clock (20). In our experiments, segmentation did not re-establish after the knock-down of two Wnt components although their expression could have resumed after blocking RNAi by the hsVSR. This indicates that the Wnt/cad feedback loop had irreversibly broken down. Interestingly, posterior Wnt signaling is self-activating at least at early stages (17). This auto regulatory loop would theoretically be sufficient to maintain Wnt activity at the posterior and the design of such a simple positive feedback loop would readily explain an irreversible breakdown. However, a simple one-component self-activating system would be prone to activation at erroneous sites – especially considering the many places of Wnt activity throughout development. Therefore, *Tc-cad* seems to be an essential component of the upstream feedback-loop maintaining the SAZ. This additional component would confer a robust localization of the SAZ at the posterior. Indeed, there is evidence that *Tc-cad* and Wnt signaling depend on each other in *T. castaneum* (15, 17, 19). Hence, we predict that in similar hsVSR experiments, *Tc-cad* RNAi would lead to an irreversible knock-down as well. Unfortunately, we were not able to test this hypothesis because strong parental RNAi targeting *Tc-cad* resulted in sterility.

The second regulatory feedback loop in the system is the primary PRG gene circuit (8). In the framework of the clock and speed regulation gradient model, this circuit is thought to be the molecular realization of the cell-autonomous clock (19). We found that this feedback loop readily re-established itself after the knock-down of the components was blocked by hsVSR treatment. This was observed for all three components. Hence, the components of that loop appear to be connected in a way that allows for a re-initiation of the clock whenever all components are functional (i.e. all three pPRGs and the upstream Wnt/cad gradient).

It had remained a possibility that an as yet unknown factor expressed in the SAZ was acting upstream of the Wnt/cad feedback loop in order to maintain the SAZ. Our data argues against such a hypothetical factor because the interruption of Wnt signaling led to an irreversible breakdown of the system. If the Wnt/cad system was activated by an upstream factor, one would have expected re-initiation. In line with this hypothesis, the genome wide iBeetle RNAi screen failed to reveal such a component in *T. castaneum* (47, 48).

### Ensuring specificity and robustness of the segmentation

The different levels of robustness of the two gene regulatory loops to external manipulations reflect the need for on one hand initiating segmentation only once and at one specific location and on the other hand ensuring the need for robustness of the ongoing segmentation towards external perturbations. Indeed, the regulatory interactions initiating segmentation seem to be designed in a way that establishment of a secondary ectopic SAZ is unlikely. At least three different signaling events need to coincide: The asymmetric activity of canonical Wnt signaling at the posterior is the first zygotic readout of maternally driven axis formation and therefore an excellent trigger for locating the SAZ posteriorly (15, 49). *Tc-tor* signaling restricts activation to very early stages because torso signaling is active in the SAZ only early on (41, 50). *Tc-cad* expression is thought to be regulated by the initial Wnt asymmetry – still, it could confer additional robustness to the spatial specificity of SAZ induction (51). Indeed, the initiation system seems to be extremely stable as we are not aware of reports of split posterior trunks in insect embryos. It would be worth testing, whether these three components are indeed sufficient to initiate a SAZ by the joint ectopic activation of these three components making use of the transgenic and genome editing tool kit of *T. castaneum* (52).

In contrast to the initiation, which should happen only once and only at one position, the ongoing segmentation process should be robust against external perturbations. Hence, our finding that the regulatory setup of the clock components allow for re-initiation after external perturbation fit that expectation. Indeed, the irregular stripes re-initiating after rescue in early embryos (Fig. 5 E) lead to remarkable well-developed segments in the cuticle. Further, when a second SAZ is specified early on either by genetic interference or by classic embryonic manipulations in other insects, a perfectly well developed mirror image abdomen can develop (15, 53–56). This indicates that indeed, after initiation, the segmentation process is robust and autonomous.

### Strains and husbandry

*Tribolium castaneum* (HERBST) beetles were reared using standard conditions and methods (52). During experiments, beetles (embryos/larvae) were kept at 32°C and 40% RH while general stock keeping was done at 28°C and 40% RH. RNAi inhibition experiments were performed in the transgenic line containing the RNAi inhibitor CrPV1A from the Cricket Paralysis Virus under the control of the endogenous *Tribolium* heat shock promoter (40) and the 3xP3DsRed (eye marker) integrated into the genome using piggyback vector (57) in *Tribolium* line vermillion^white^. Non-transgenic vermillion^white^ beetles were used for negative controls.

### RNAi and heat-shock treatment

Parental RNAi was performed according to established methods (5, 58). Templates were prepared by PCR with T7 overhanging-primers from plasmid templates containing varying lengths of coding and/or regulatory mRNA sequence (*Tc-even-skipped* ∼1400 bp, *Tc-odd-skipped* ∼380 bp, *Tc-paired* ∼540 bp, *Tc-arrow* ∼1800 bp, *Tc-Wnt8/D* ∼500 bp, *Tc-wntless* ∼600 bp). DsRNA was produced using MEGAscript T7 Transcription Kit (Life Technologies). The concentration of injected dsRNA for parental RNAi was 1000 ng/μl (*Tc-even-skipped*), 500 ng/μl (*Tc-odd-skipped, Tc-paired*), or 100 ng/μl (*Tc-arrow, Tc-Wnt8/D, Tc-wntless*). Add NCBI accession number? *Tc-eve*: NM_001039449.1; *Tc-odd*: XM_008198532.2; *Tc-run*: XM_964184.3; *Tc-wg*: NM_001114350

Embryos collected from dsRNA injected animals of either the hsVSR or wild type control (*vermillion^whit^*^e^) were collected and kept without flour at 32°C in small plastic fly culturing vials before the treatment. For the heat-shock, the eggs were transferred to a small (40ml) glass beaker with a flat bottom making sure that all embryos had direct contact with the bottom. Then, the beaker was put into a pre-heated 48°C waterbath. The beaker was covered with perforated aluminum foil. The bottom of the glass beaker was kept submerged for 10 min. To ensure a controlled and quick termination of the heatshock, the beaker was put into a room temperature waterbath. After transferring the embryos back into the plastic vials, they were allowed to recover for two hours at 32°C, until they were heatshocked a second time for 10 min at 48°C, following the same procedure. Thereafter, the embryos were kept at 32°C until fixation or cuticle preparation. We note that the results were sensitive even to minor changes of the procedure and each step needed to be optimized carefully.

### Staining, and microscopy

Embryo fixation and in-situ hybridisation were performed as described previously (59). Digoxigenin (DIG)-labeled riboprobes targeting *Tc-wg* (DIG RNA Labeling Kit, Roche), was detected by anti-DIG-AP antibodies (Roche) and visualized by NBT/BCIP staining. HCR staining was performed as published with small modification (kindly provided by Eric Clark and Olivia Tidswell prior publication) (60, 61). HCR probes for *Tc-eve* were purchased from Molecular Technologies while HCR probes for all other genes were purchased from Molecular Instruments. Binding sequences are available from vendors at request due to intellectual properties restrictions. Cuticles of L1 larvae were analyzed and documented using either a Zeiss AxioPlan 2 (10x air objective) with ImagePro 6 or a Leica SP5 inverted cLSM (10x air objective) with Leica LAS-X software, utilizing the cuticle’s autofluorescence. HCR stainings were documented using a Leica SP8 confocal laser-scanning microscope (20x objectives with 100% glycerol as immersion medium) and the Leica LAS-X software (v 3.5.2). In-situ hybridization was documented using Zeiss AxioPlan 2 or Zeiss AxioScope.

### qPCR

RNA from whole embryos was extracted using the Quick-RNA Tissue/Insect Kit (Zymo Research) with DNase on-column digest (DNaseI Set, Zymo Research). cDNA was synthesized using the MAXIMA First Strand cDNA Synthesis Kit for RT-qPCR (Thermo Fisher Scientific) according to manufacturer’s instructions. qPCRs were performed using the CFX96 Real-Time PCR System (Bio-Rad Laboratories) with 5x HOT FIREPol® EvaGreen® qPCR Mix Plus (ROX) Master mix (Solis Biodyne). Reference genes were identified using RefFinder (62). qPCR data analysis was done in the CFX Manager 3.1 (Bio-Rad Laboratories) and pyQPCR with the delta-delta-Ct method (63).

### Statistical analysis

Comparisons of abdominal segment numbers in cuticles and comparisons of number of expression stripes in germbands were tested using unpaired, two-sided Mann–Whitney U tests for independent samples. All measured data points were included in the calculations and were not checked for outliers beforehand. Outliers were determined for the plots using the R package *ggplot2*, considering data above 1.5 *IQR of the 75th percentile or below 1.5 *IQR of the 25th percentile as outliers, which are indicated in in the respective plots in red. Comparisons of the number of stripes in germbands were done using the Pearson’s Chi-squared Test for Count Data with simulated p-values by Monte Carlo simulations (B=1000). All graphs and statistical calculations were performed using R (v3.5.2; R Core Team, 2018) and RStudio (v1.1.x; RStudio Team, 2015) with the following packages: dplyr, ggplot2, ggpubr, ggsigni, patchwork, readxl, reshape2.

## Acknowledgements

We thank Michalis Averof and Martin Klingler for valuable discussions and Eric Clark, Olivia Tidswell and Michael Akam for sharing protocols prior publication. This work was funded by Deutsche Forschungsgemeinschaft ANR-DFG joint programme (together with M. Averof) BU1443/14-1 and the Deutsche Forschungsgemeinschaft programme for researchers in danger BU1443/14-1ad.

## References

1. E. Clark, A. D. Peel, M. Akam, Arthropod segmentation. Development (Cambridge, England) 146, dev170480 (2019).

2. G. K. Davis, N. H. Patel, SHORT, LONG, AND BEYOND: Molecular and Embryological Approaches to Insect Segmentation. Annual Review of Entomology 47, 669–699 (2002).

3. D. Tautz, Segmentation. Dev Cell 7, 301–12 (2004).

4. M. Diaz-Cuadros, O. Pourquié, E. El-Sherif, Patterning with clocks and genetic cascades: Segmentation and regionalization of vertebrate versus insect body plans. PLoS Genet 17, e1009812 (2021).

5. G. Bucher, J. Scholten, M. Klingler, Parental RNAi in Tribolium (Coleoptera). Current Biology 12, R85–R86 (2002).

6. C. Nüsslein-Volhard, E. Wieschaus, Mutations affecting segment number and polarity in Drosophila. Nature 287, 795–801 (1980).

7. D. St Johnston, C. Nüsslein-Volhard, The origin of pattern and polarity in the Drosophila embryo. Cell 68, 201–19 (1992).

8. C. P. Choe, S. C. Miller, S. J. Brown, A pair-rule gene circuit defines segments sequentially in the short-germ insect Tribolium castaneum. Proceedings of the National Academy of Sciences of the United States of America 103, 6560–6564 (2006).

9. E. Clark, Dynamic patterning by the Drosophila pair-rule network reconciles long-germ and short-germ segmentation. PLoS Biol 15, e2002439 (2017).

10. E. El-Sherif, M. Averof, S. J. Brown, A segmentation clock operating in blastoderm and germband stages of Tribolium development. *Development (Cambridge*, England) 139, 4341– 4346 (2012).

11. A. F. Sarrazin, A. D. Peel, M. Averof, A segmentation clock with two-segment periodicity in insects. Science 336, 338–341 (2012).

12. A. C. Oates, L. G. Morelli, S. Ares, Patterning embryos with oscillations: structure, function and dynamics of the vertebrate segmentation clock. Development 139, 625–639 (2012).

13. I. Palmeirim, D. Henrique, D. Ish-Horowicz, O. Pourquié, Avian hairy gene expression identifies a molecular clock linked to vertebrate segmentation and somitogenesis. Cell 91, 639– 648 (1997).

14. H. Rudolf, C. Zellner, E. El-Sherif, Speeding up anterior-posterior patterning of insects by differential initialization of the gap gene cascade. Dev Biol 460, 20–31 (2020).

15. S. Ansari, et al., Double abdomen in a short-germ insect: Zygotic control of axis formation revealed in the beetle Tribolium castaneum. Proc. Natl. Acad. Sci. U.S.A. 115, 1819– 1824 (2018).

16. A. Beermann, R. Pruhs, R. Lutz, R. Schroder, A context-dependent combination of Wnt receptors controls axis elongation and leg development in a short germ insect. Development 138, 2793–805 (2011).

17. G. Oberhofer, D. Grossmann, J. L. Siemanowski, T. Beissbarth, G. Bucher, Wnt/β- catenin signaling integrates patterning and metabolism of the insect growth zone. Development, dev.112797 (2014).

18. A. Schönauer, et al., The Wnt and Delta-Notch signalling pathways interact to direct pair-rule gene expression via caudal during segment addition in the spider Parasteatoda tepidariorum. Development 143, 2455–2463 (2016).

19. E. El-Sherif, X. Zhu, J. Fu, S. J. Brown, Caudal Regulates the Spatiotemporal Dynamics of Pair-Rule Waves in Tribolium. PLOS Genetics 10, e1004677 (2014).

20. X. Zhu, et al., Speed regulation of genetic cascades allows for evolvability in the body plan specification of insects. Proceedings of the National Academy of Sciences of the United States of America 128, 201702478–E8655 (2017).

21. E. Clark, A. D. Peel, Evidence for the temporal regulation of insect segmentation by a conserved sequence of transcription factors. Development 145, dev155580, dev.155580 (2018).

22. M. Aranda, H. Marques-Souza, T. Bayer, D. Tautz, The role of the segmentation gene hairy in Tribolium. Dev. Genes Evol. 218, 465–477 (2008).

23. C. P. Choe, S. J. Brown, Evolutionary flexibility of pair-rule patterning revealed by functional analysis of secondary pair-rule genes, paired and sloppy-paired in the short-germ insect, Tribolium castaneum. Developmental biology 302, 281–294 (2007).

24. F. Maderspacher, G. Bucher, M. Klingler, Pair-rule and gap gene mutants in the flour beetle Tribolium castaneum. Dev Genes Evol 208, 558–68 (1998).

25. A. Boos, J. Distler, H. Rudolf, M. Klingler, E. El-Sherif, A re-inducible gap gene cascade patterns the anterior-posterior axis of insects in a threshold-free fashion. Elife 7, e41208 (2018).

26. G. Bucher, M. Klingler, Divergent segmentation mechanism in the short germ insect Tribolium revealed by giant expression and function. Development 131, 1729–40 (2004).

27. A. C. Cerny, G. Bucher, R. Schroder, M. Klingler, Breakdown of abdominal patterning in the Tribolium Kruppel mutant jaws. *Development (Cambridge*, England) 132, 5353–63 (2005).

28. H. Marques-Souza, M. Aranda, D. Tautz, Delimiting the conserved features of hunchback function for the trunk organization of insects. *Development (Cambridge*, England) 135, 881–8 (2008).

29. R. Aliyari, et al., Mechanism of induction and suppression of antiviral immunity directed by virus-derived small RNAs in Drosophila. Cell Host Microbe 4, 387–397 (2008).

30. E. H. Bayne, D. V. Rakitina, S. Y. Morozov, D. C. Baulcombe, Cell-to-cell movement of potato potexvirus X is dependent on suppression of RNA silencing. Plant J 44, 471–482 (2005).

31. J. A. Chao, et al., Dual modes of RNA-silencing suppression by Flock House virus protein B2. Nat Struct Mol Biol 12, 952–957 (2005).

32. H. Jin, J.-K. Zhu, A viral suppressor protein inhibits host RNA silencing by hooking up with Argonautes. Genes Dev 24, 853–856 (2010).

33. A. Nayak, et al., Cricket paralysis virus antagonizes Argonaute 2 to modulate antiviral defense in Drosophila. Nat Struct Mol Biol 17, 547–554 (2010).

34. J. T. van Mierlo, et al., Convergent evolution of argonaute-2 slicer antagonism in two distinct insect RNA viruses. PLoS Pathog 8, e1002872 (2012).

35. R. P. van Rij, et al., The RNA silencing endonuclease Argonaute 2 mediates specific antiviral immunity in Drosophila melanogaster. Genes Dev 20, 2985–2995 (2006).

36. O. Voinnet, C. Lederer, D. C. Baulcombe, A viral movement protein prevents spread of the gene silencing signal in Nicotiana benthamiana. Cell 103, 157–167 (2000).

37. G. Meister, T. Tuschl, Mechanisms of gene silencing by double-stranded RNA. Nature 431, 343–9 (2004).

38. C. B. Phelps, A. H. Brand, Ectopic gene expression in Drosophila using GAL4 system. Methods 14, 367–79 (1998).

39. J. B. Schinko, et al., Functionality of the GAL4/UAS system in Tribolium requires the use of endogenous core promoters. BMC Dev Biol 10, 53 (2010).

40. J. Schinko, K. Hillebrand, G. Bucher, Heat shock-mediated misexpression of genes in the beetle Tribolium castaneum. Dev Genes Evol. **Dev Genes Evol.**, 287–98 (2012).

41. M. Schoppmeier, R. Schröder, Maternal torso signaling controls body axis elongation in a short germ insect. Current Biology 15, 2131–2136 (2005).

42. R. Bolognesi, L. Farzana, T. D. Fischer, S. J. Brown, Multiple wnt genes are required for segmentation in the short-germ embryo of tribolium castaneum. Current biology : CB 18, 1624– 1629 (2008).

43. T. Copf, R. Schroder, M. Averof, Ancestral role of caudal genes in axis elongation and segmentation. Proceedings of the National Academy of Sciences of the United States of America 101, 17711–5 (2004).

44. E. V. W. Setton, P. P. Sharma, A conserved role for arrow in posterior axis patterning across Arthropoda. Dev Biol 475, 91–105 (2021).

45. R. Bolognesi, T. D. Fischer, S. J. Brown, Loss of Tc-arrow and canonical Wnt signaling alters posterior morphology and pair-rule gene expression in the short-germ insect, Tribolium castaneum. Development Genes and Evolution 219, 369–375 (2009).

46. R. Bolognesi, et al., Tribolium Wnts: evidence for a larger repertoire in insects with overlapping expression patterns that suggest multiple redundant functions in embryogenesis. Development genes and evolution 218, 193–202 (2008).

47. M. S. Hakeemi, et al., Screens in fly and beetle reveal vastly divergent gene sets required for developmental processes. BMC Biology 20, 38 (2022).

48. C. Schmitt-Engel, et al., The iBeetle large-scale RNAi screen reveals gene functions for insect development and physiology. Nature Communications 6, 7822 (2015).

49. J. Fu, et al., Asymmetrically expressed axin required for anterior development in Tribolium. Proc. Natl. Acad. Sci. U.S.A. 109, 7782–7786 (2012).

50. R. Schröder, C. Eckert, C. Wolff, D. Tautz, Conserved and divergent aspects of terminal patterning in the beetle Tribolium castaneum. Proc. Natl. Acad. Sci. U.S.A. 97, 6591– 6596 (2000).

51. C. Schulz, R. Schroder, B. Hausdorf, C. Wolff, D. Tautz, A caudal homologue in the short germ band beetle Tribolium shows similarities to both, the Drosophila and the vertebrate caudal expression patterns. Development genes and evolution 208, 283–9 (1998).

52. M. Klingler, G. Bucher, The red flour beetle T. castaneum: elaborate genetic toolkit and unbiased large scale RNAi screening to study insect biology and evolution. Evodevo 13, 14 (2022).

53. K. Sander, [Reversal of the germ band polarity in egg fragments of Euscelis (Cicadina)]. Experientia 17, 179–180 (1961).

54. K. Sander, “Specification of the basic body pattern in insect embryogenesis” in Advances in Insect Physology, J. E. Treherne, M. J. Berridge, V. B. Wigglesworth, Eds. (Academic Press, 1976), pp. 125–238.

55. J. Klomp, et al., Embryo development. A cysteine-clamp gene drives embryo polarity in the midge Chironomus. Science (New York, N.Y.) 348, 1040–1042 (2015).

56. Y. Yoon, et al., Embryo polarity in moth flies and mosquitoes relies on distinct old genes with localized transcript isoforms. Elife 8, e46711 (2019).

57. A. J. Berghammer, M. Klingler, E. A. Wimmer, A universal marker for transgenic insects. Nature 402, 370–1 (1999).

58. N. Posnien, et al., RNAi in the red flour beetle (Tribolium). Cold Spring Harb Protoc 2009, pdb.prot5256 (2009).

59. J. Schinko, N. Posnien, S. Kittelmann, N. Koniszewski, G. Bucher, Single and double whole-mount in situ hybridization in red flour beetle (Tribolium) embryos. Cold Spring Harb Protoc 2009, pdb.prot5258 (2009).

60. H. M. T. Choi, et al., Third-generation in situ hybridization chain reaction: multiplexed, quantitative, sensitive, versatile, robust. Development 145, dev165753 (2018).

61. O. R. A. Tidswell, M. A. Benton, M. Akam, The neuroblast timer gene nubbin exhibits functional redundancy with gap genes to regulate segment identity in Tribolium. Development 148, dev199719 (2021).

62. F. Xie, J. Wang, B. Zhang, RefFinder: a web-based tool for comprehensively analyzing and identifying reference genes. Funct Integr Genomics 23, 125 (2023).

63. T. D. Schmittgen, K. J. Livak, Analyzing real-time PCR data by the comparative CT method. Nat. Protocols 3, 1101–1108 (2008).

