## Supplementary data for "Temporal restriction of RNAi reveals breakdown of the segmentation clock is reversible after knock down of primary pair rule genes but not Wnt-signaling in the red flour beetle"

### Supplementary Text

#### 1) Establishment of CrPV1A as a tool for suppressing RNAi in *T. castaneum*

Six VSRs were tested in order to identify a strong Viral Suppressor of RNAi (VSR) in *T. castaneum*. The inhibitors from the *Cricket Paralysis virus* (CrPV1A), the *Flock House virus* (FHV B2), *Drosophila C virus* (DCV1A) and *Nora virus* (VP1) had previously been shown to function as suppressors of the RNAi mechanism in *Drosophila melanogaster* (1–6). Further, the suppressor proteins p25 from the Potato virus X (PVX) and p38 from the Turnip Crinkle virus (TCV) were tested. These plant virus proteins had been known to inhibit the RNAi pathway, but not the microRNA pathway in plants (7–9).

First, we evaluated if any of those VSRs would be able to block RNAi targeting an exogenous gene expressed from a transgenic construct. For this purpose, we crossed a UAS-tGFP responder line (10) (red eye marker) to a driver line that expressed Gal4delta in a thoracic region during late larval, pupal and adult stages (name: *Bauchbinde-Gal4*; black eye marker). The double heterozygous offspring of this mating showed a strong thoracic fluorescence (Fig. A – leftmost columns). In parallel, we generated animals where in addition to UAS-tGFP also the UAS-VSR construct was present (right columns). We performed RNAi targeting tGFP both, in the cross without UAS-VSR (middle column) and with UAS-VSR (right column; see table for overview). The RNAi was predicted to deplete tGFP expression unless the UAS-VSR was able to block RNAi. In this experiment, only CrPV1A blocked RNAi to a degree that tGFP expression remained visible (Fig. A top left panel, red box).

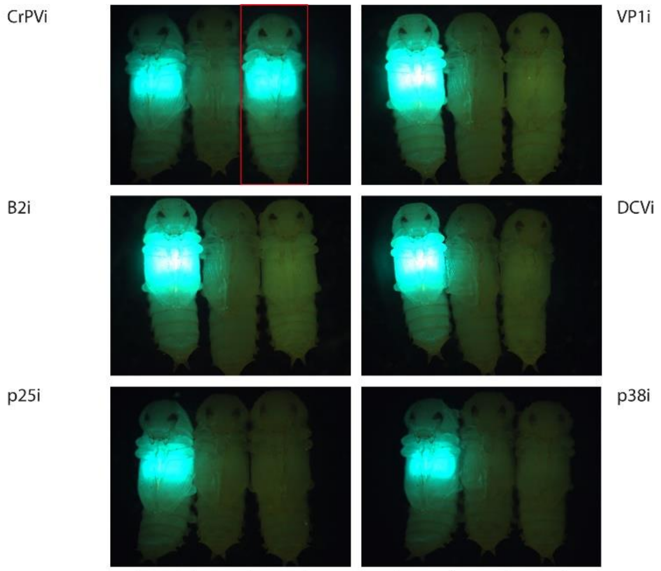

| Treatment | 1 | 2 | 3 | | 1 | 2 | 3 |
|---|---|---|---|---|---|---|---|
| BB-Gal4 X UAS-tGFP | X | X | X | | X | X | X |
| BB-Gal4 X UAS-tGFP X UAS-VSR | | | X | | | | X |
| tGFP RNAi | | X | X | | | X | X |

**Figure A Testing six VSRs for inhibition of RNAi**. See text for details

In a second experiment, we tested the VSR effect with respect to RNAi targeting an endogenous gene. Larval RNAi against the pigmentation gene *Tc-ebony* was performed. This gene is an N-beta-alanyl dopamine synthetase that is necessary for the synthesis of NBAD sclerotin. RNAi knocking down *Tc-ebony* leads to darkened or black body pigmentation. In this experiment, the UAS-VSR responder lines were crossed to a Gal4 driver, which showed ubiquitous Gal4delta-expression in the epidermis except for early embryonic stages (until appr. 24h at 32°C; not shown; we did not check internal organs; name: *Boje-Gal4*). In the presence of an active VSRs a reduction of the portion of black animals and an increase of animals with wildtype color was expected. Again, CrVP1A showed a complete rescue for one insertion (Fig. B, panel D) and partial rescue for another insertion (Fig. B, panel C). The FHV B2 VSR was active to a lesser degree (Fig. B panels E,F - two different insertions led to an increase of wt coloured animals).

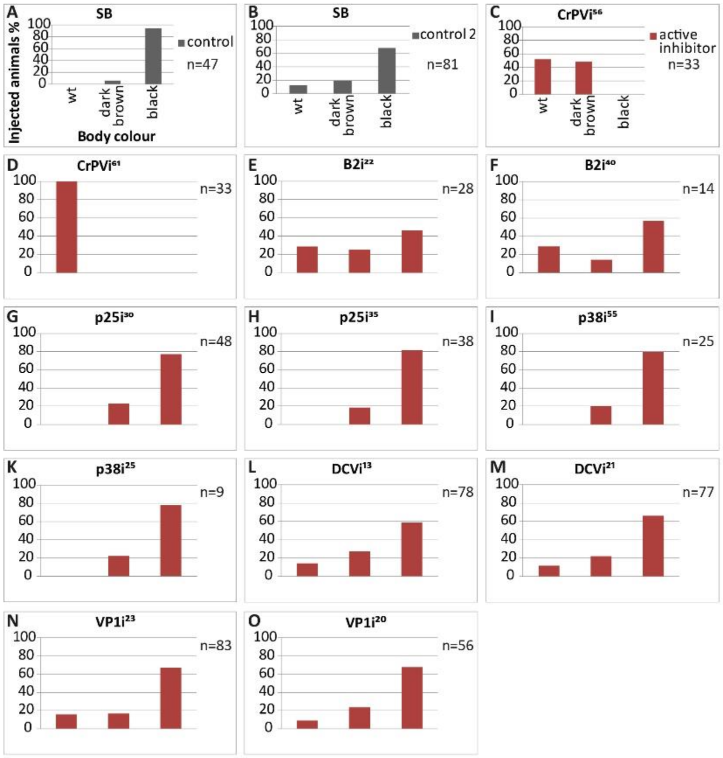

**Figure B Testing six VSRs for inhibition of RNAi**
See text for details

Taken together, these tests revealed CrPV1A as an efficient suppressor of RNAi in *T. castaneum* transgenically expressed. Based on these results, we chose CrPV1A as tool for temporal control of RNAi in *T. castaneum*.

#### 2) The VSR CrPV1A

*Cricket Paralysis virus* (CrPV) was initially identified and isolated from field crickets, *Teleogryllus oceanicus* and *Teleogryllus commodus,* and it is a highly potent virus of many species in the laboratory (4, 11, 11). CrPV is closely related to *Drosophila C virus* and likewise belongs to the positive-strand *Dicistroviridae* family. In contrast to DCV, CrPV leads to mortality upon infection of crickets and flies (4, 12). This high pathogenesis of CrPV is partially based on its efficient RNAi suppressor protein, CrPV1A. It has been shown that adding CrPV1A to the Sindbis virus, which does not naturally encode an endogenous RNAi suppressor, resulted in increased virus production and fly lethality upon infection (4). The mode of action of CrPV1A relies on its interaction with the endonuclease Ago-2, a component of the RISC complex. This interaction blocks Ago-2 cleavage activity, resulting in inhibited RISC-mediated mRNA degradation and therewith RNAi disruption. Nevertheless, the suppressor protein CrPV1A did not interfere with the miRNA pathway or alter the physiology and development of the animals when expressed in flies (4). Note that our results indicate that overexpression of CrPV1A interferes with viability, possibly via blocking the miRNA pathway.

### Supplementary Figures

#### Fig. S1 Embryonic schemes and replicate of *Tc-paired* experiments

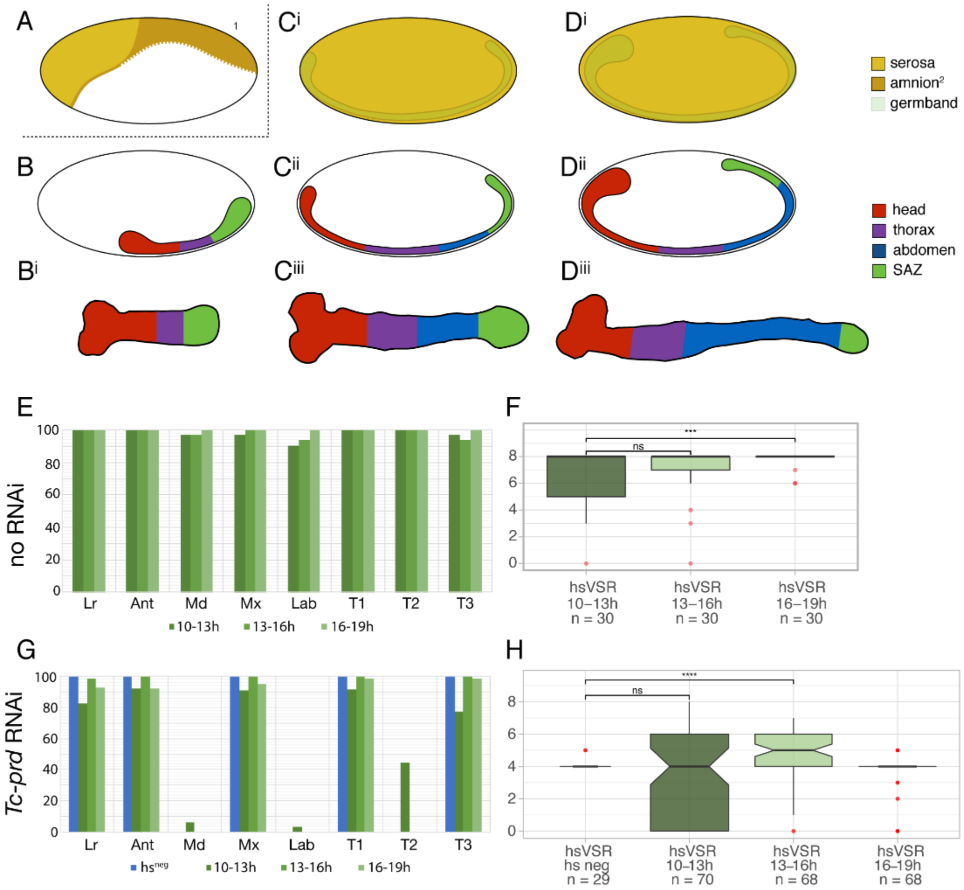

**Fig. S1 Scheme of early development and repetition of control experiments**

A) In *T. castaneum*, the embryonic anlagen are located at a ventral posterior position in the blastoderm. The extraembryonic tissues serosa (light yellow) and amnion (dark yellow) are depicted. B) The early germ band can be separated in head and thoracic segments (red and purple), which are specified during the blastoderm stage and the SAZ (green). C,D) During elongation, abdominal segments are added in a sequential way in the SAZ. The depicted stages correspond approximately to the stages in which heat-shocks were performed. E) Heat-shocks alone did not much affect anterior segments. F) Importantly, the earlier heat-shocks did affect the number of abdominal segments indicating that this treatment disturbs segmentation to some degree. G and H) Repetition of the Tc-prd experiment (see text for details). In this repetition, the negative influence of the heat-shock is more pronounced than in the experiment shown in Fig. 2.

#### Fig. S2 Repetition of Wnt-signalling experiments

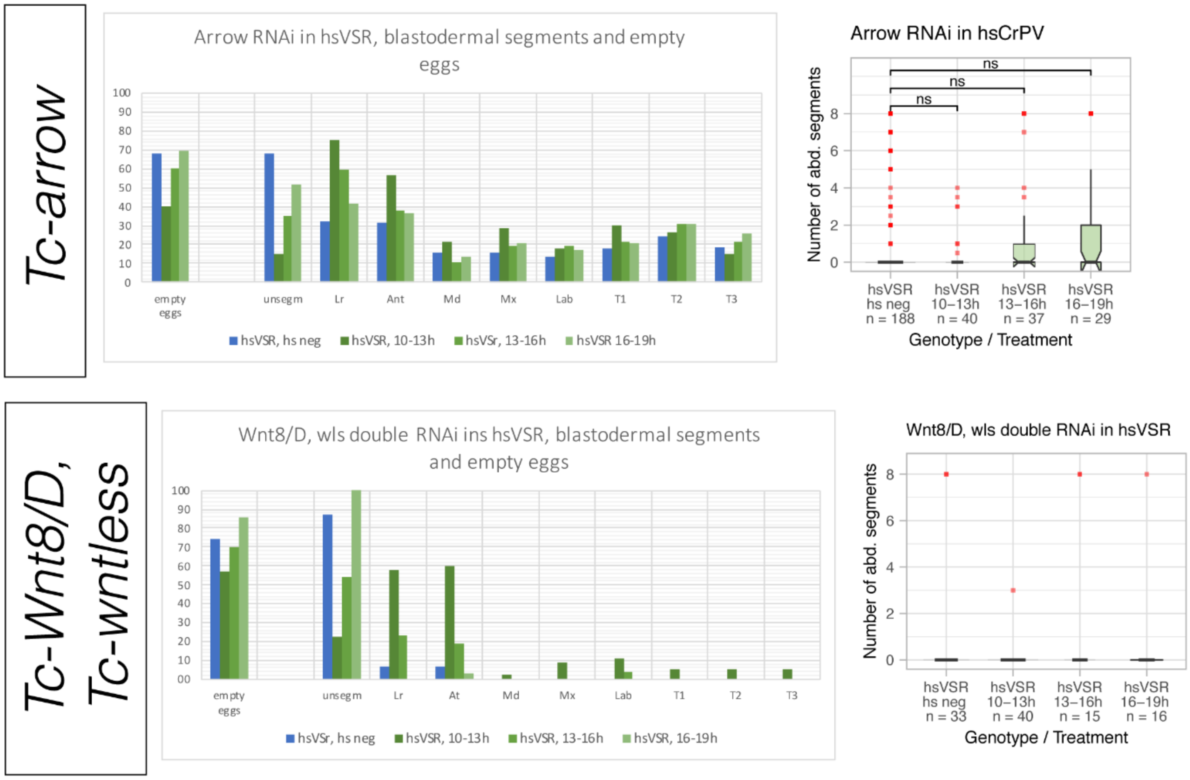

### Fig. S3 Repetition of Tc-even-skipped experiment

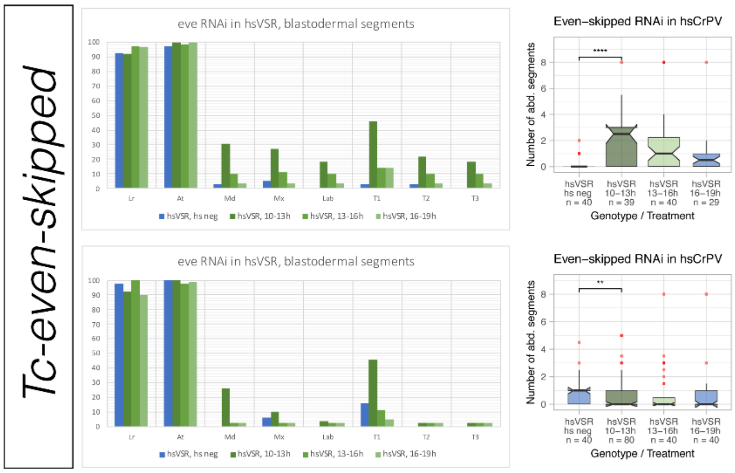

#### Fig. S4 Replicates by independent researcher

Controls:

*Tc-paired*

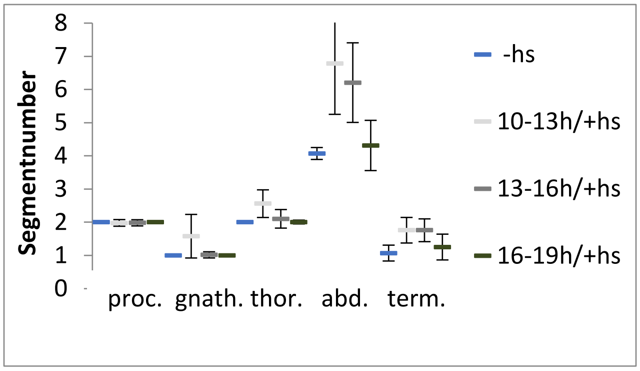

*wildtype*

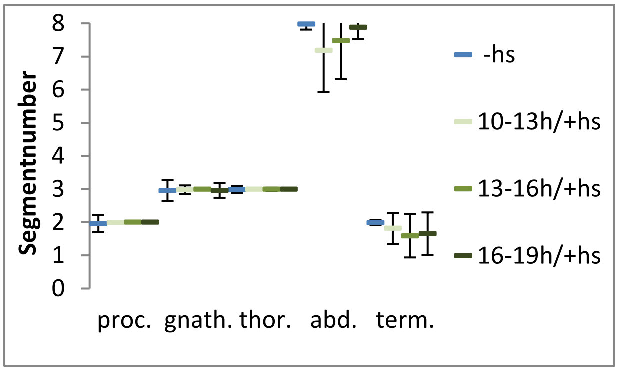

*Tc-odd*

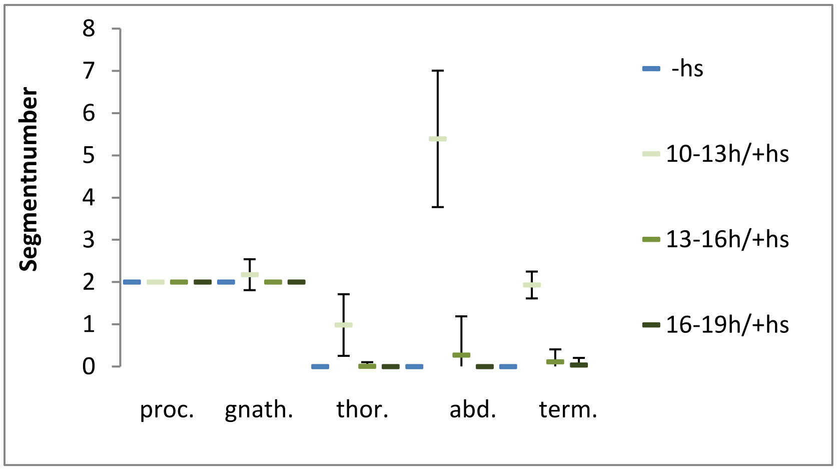

*Tc-runt*

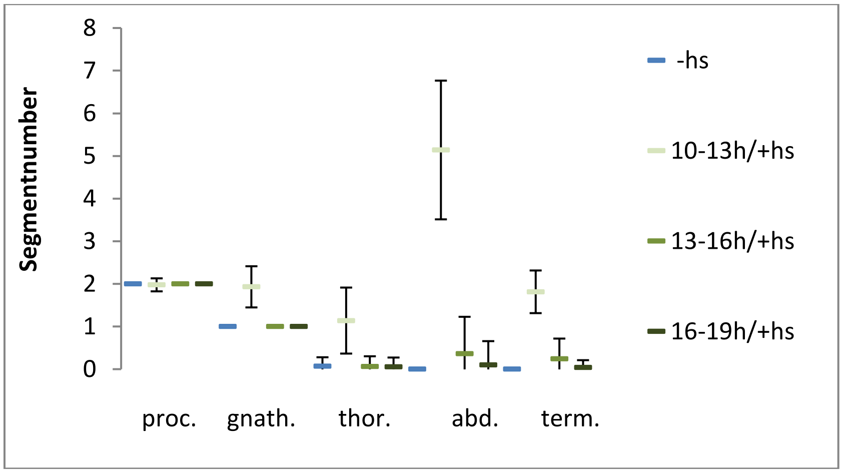

#### Fig. S5 Definition of developmental stages based on head expression of *Tc-wg*

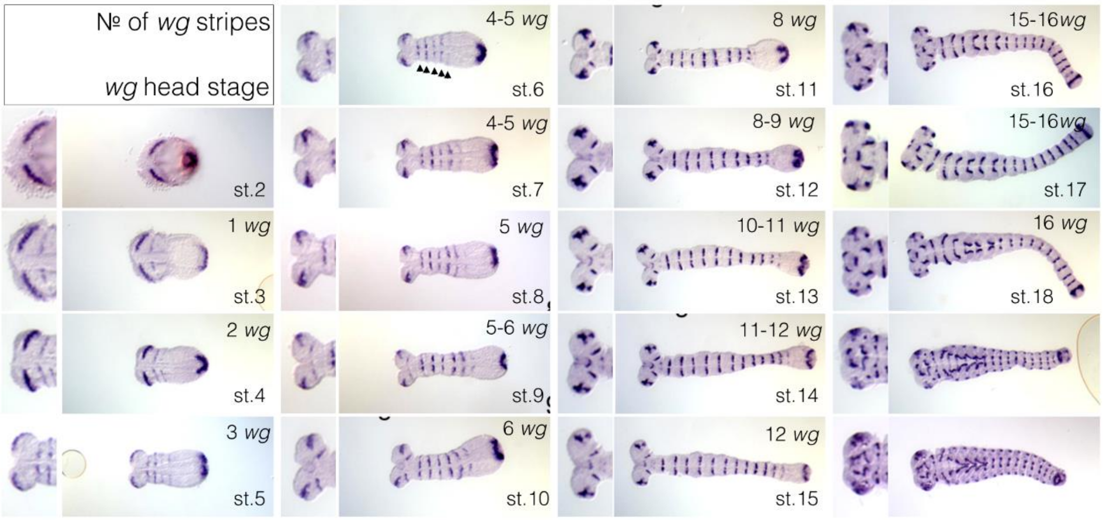

#### Fig. S6 Determining the developmental delay induced by heat-shock

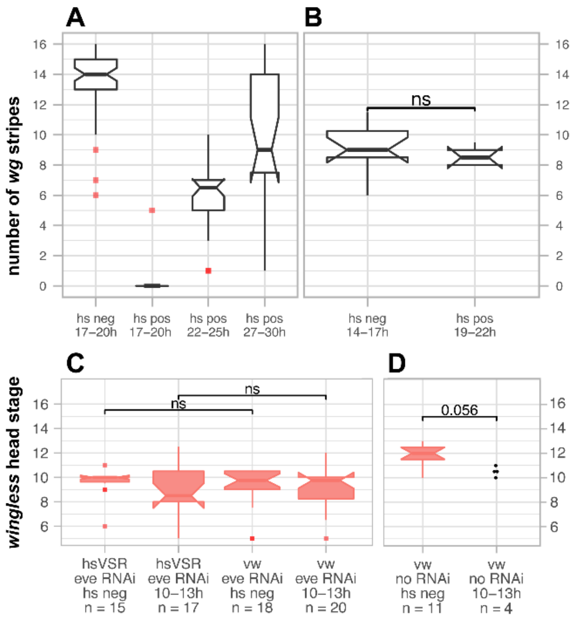

A) To determine the delay, we first compared development between wt and heat-shocked embryos using the number of abdominal *Tc-wg* stripes. For late stages, this revealed a lag of more than 10 hours. B) For embryos during ongoing segmentation, fixation with a 5h difference resulted in comparable stages. C) We used that timing to stage heat-shocked and non-heat-shocked wt and *Tc-eve*-RNAi embryos. Because these RNAi embryos miss abdominal segments, we used the dynamic *Tc-wg* head expression to stage the embryos (see Fig. S5). D) The shift of developmental timing was confirmed by counting Tc-wg stripes in embryos without RNAi.

#### Fig. S7 Rescued Tc-eve embryos and classification

See text for details

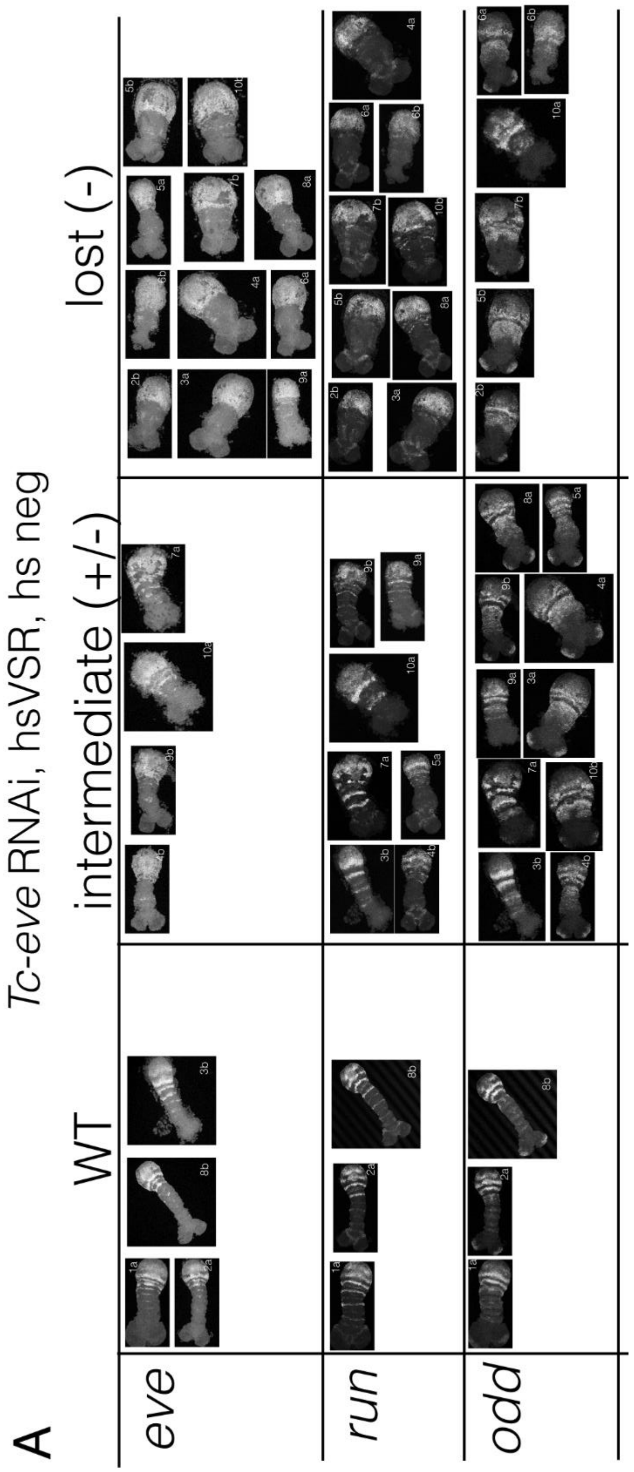

#### Fig. S8 Measuring the knock-down by qPCR

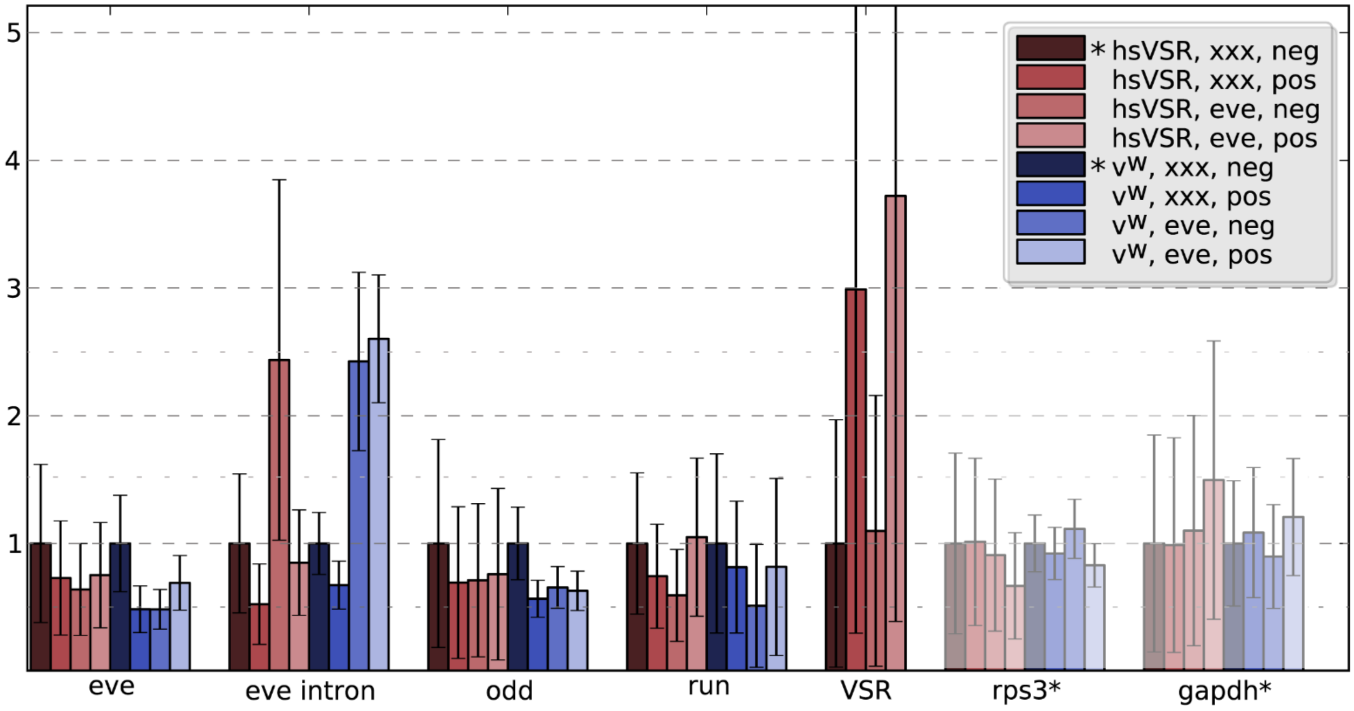

qPCR was performed in the hsVSR line (reddish colors) and vw wildtype (blue colors). Rps3 and gapdh were used for normalization. As expected, heat-shock treatment increased the amount of VSR transcript (VSR – compare bars with heatshock (“pos”) with those without heatshock (“neg”). Surprisingly, Tc-eve expression was not much reduced after Tc-eve RNAi. However, nuclear transcripts did increase dramatically in eve RNAi embryos that were not rescued (eve intron, intermediate red bar). Tc-odd expression was not much altered in line with ongoing expression in the knock-down embryos. Tc-runt  may be downregulated in Tc-eve RNAi (middle bar) but the expression was rescued in Tc-eve RNAi combined with hsVSR rescue (light red bar).

#### References

1. R. Aliyari, *et al.*, Mechanism of induction and suppression of antiviral immunity directed by virus-derived small RNAs in Drosophila. *Cell Host Microbe* **4**, 387–397 (2008).

2. J. A. Chao, *et al.*, Dual modes of RNA-silencing suppression by Flock House virus protein B2. *Nat Struct Mol Biol* **12**, 952–957 (2005).

3. H. Li, W. X. Li, S. W. Ding, Induction and suppression of RNA silencing by an animal virus. *Science* **296**, 1319–1321 (2002).

4. A. Nayak, *et al.*, Cricket paralysis virus antagonizes Argonaute 2 to modulate antiviral defense in Drosophila. *Nat Struct Mol Biol* **17**, 547–554 (2010).

5. J. T. van Mierlo, *et al.*, Convergent evolution of argonaute-2 slicer antagonism in two distinct insect RNA viruses. *PLoS Pathog* **8**, e1002872 (2012).

6. R. P. van Rij, *et al.*, The RNA silencing endonuclease Argonaute 2 mediates specific antiviral immunity in Drosophila melanogaster. *Genes Dev* **20**, 2985–2995 (2006).

7. E. H. Bayne, D. V. Rakitina, S. Y. Morozov, D. C. Baulcombe, Cell-to-cell movement of potato potexvirus X is dependent on suppression of RNA silencing. *Plant J* **44**, 471–482 (2005).

8. H. Jin, J.-K. Zhu, A viral suppressor protein inhibits host RNA silencing by hooking up with Argonautes. *Genes Dev* **24**, 853–856 (2010).

9. O. Voinnet, C. Lederer, D. C. Baulcombe, A viral movement protein prevents spread of the gene silencing signal in Nicotiana benthamiana. *Cell* **103**, 157–167 (2000).

10. J. B. Schinko, *et al.*, Functionality of the GAL4/UAS system in *T. castaneum* requires the use of endogenous core promoters. *BMC Dev Biol* **10**, 53 (2010).

11. N. Plus, G. Croizier, C. Reinganum, P. D. Scott, Cricket paralysis virus and drosophila C virus: serological analysis and comparison of capsid polypeptides and host range. *J Invertebr Pathol* **31**, 296–302 (1978).

12. T. Manousis, N. F. Moore, Cricket Paralysis Virus, a Potential Control Agent for the Olive Fruit Fly, Dacus oleae Gmel. *Appl Environ Microbiol* **53**, 142–148 (1987).
